# gSV: a general structural variant detector using the third-generation sequencing data

**DOI:** 10.64898/2026.03.02.703663

**Authors:** Jingyu Hao, Jiandong Shi, Sheng Lian, Zhen Zhang, Yongyi Luo, Taobo Hu, Toyotaka Ishibashi, Depeng Wang, Shu Wang, Xiaodan Fan, Weichuan Yu

## Abstract

Structural variants (SVs) are major contributors to genome diversity and disease susceptibility, particularly in cancer. Although third-generation sequencing technologies have substantially improved SV detection sensitivity, accurate detection of complex SVs remains challenging due to fragmented and heterogeneous alignment signals, as well as the dependence of many existing methods on predefined variant models. In this paper, we propose gSV, a general SV detector that integrates alignment-based and assembly-based approaches with the maximum exact match (MEM) strategy, with particular emphasis on resolving SVs with complex or atypical alignment signatures. Without predefined assumptions about SV types, gSV captures diverse variant signals, enabling the detection of SVs that are usually missed by conventional tools. Benchmarking using both simulated datasets and real long-read sequencing data demonstrates that gSV achieves improved sensitivity and overall detection performance compared with current state-of-the-art SV callers, particularly for simple and complex SV events with complex alignment patterns. Unique SV discoveries in four breast cancer cell lines, particularly in cancer-associated genes, demonstrate the potential biological relevance of gSV-enabled discoveries. Furthermore, analysis of a breast cancer cohort from the Chinese population highlights the utility of gSV for population-scale genomic studies. Collectively, gSV provides a unified framework for comprehensive SV discovery in both research and clinical genomics settings.

##### Key Points

- Existing structural variant (SV) detection tools are limited in resolving SVs with complex alignment patterns due to their reliance on predefined variant models.
- gSV integrates alignment-based and assembly-based evidence using a maximum exact match (MEM) strategy, enabling capture of diverse and complex SV signals.
- Benchmarking on simulated and real long-read sequencing datasets demonstrates that gSV achieves competitive performance on canonical SV classes and improved sensitivity for complex SV patterns.
- Application of gSV to breast cancer cell lines and a population-scale breast cancer cohort reveals previously unresolved SVs in cancer-associated genes, highlighting its utility in genomic and clinical studies.

## Introduction

The advent of the third-generation sequencing technologies, such as PacBio and Nanopore, has enabled the acquisition of longer reads spanning thousands of base pairs (bps). Consequently, longer genomic mutations such as structural variants (SVs) are of increasing interest. SVs may be of different levels of complexity. Simple SVs can be categorized as balanced (e.g., inversions (INVs)) or unbalanced (e.g., deletions(DELs), insertions(INSs), and duplications(DUPs)). Complex SVs are usually regarded as combinations of simple SVs in various orders and numbers. These events, which often involve multiple breakpoints, nested rearrangements, or complex alignment patterns that are difficult to resolve using standard SV representations, have been widely observed in real datasets and are of increasing research interest [4]. This offers a new angle to study genetic abnormalities associated with diseases.

Many tools have been developed to detect SVs. Most of them focus on detecting simple SVs. They can be broadly categorized into alignment-based and assembly-based methods. Alignment-based methods typically involve three steps: capturing SV signatures from alignment results; clustering the signatures in terms of positions, lengths, and types; and finalizing the clustered signatures to call SVs and filtering using quality metrics (e.g., supporting reads) [21; 24; 9; 11; 3]. Alignment-based methods are popular due to their simplicity and high sensitivity, particularly in detecting simple SVs. However, existing methods can only detect potential SVs that conform to their predefined patterns at the initial step, thereby limiting their ability to resolve complex SVs with novel structural patterns that deviate from these predefined categories. Assembly-based methods, on the other hand, first assemble reads into longer consensus and subsequently align the consensus to the reference genome to detect SVs [30]. Assembly-based methods are computationally expensive and are usually not primarily optimized for resolving complex SVs. Recently, Denti et al. [6] have proposed to use both alignment-based and assembly-based methods. But their method can only detect INSs and DELs. Lin et al. [15] have applied a deep learning method to detect both simple and complex SVs, and Wang et al. [28] have further improved it. However, the issues of overfitting and weak interpretability are not tackled when applying the deep learning method. To date, accurately detecting both simple and complex SVs in an interpretable manner remains challenging, particularly for complex SVs where achieving satisfactory sensitivity and specificity (recall and precision)is still difficult.

In this paper, we propose a general SV detection framework, gSV, which integrates alignment-based and assembly-based strategies with a maximum exact match (MEM), with a particular focus on improving the detection of complex SVs. Unlike the most widely used SV detection tools that rely on prior assumptions about the types of SVs, gSV comprehensively captures a broad range suspicious variant signals. This enables the detection of SVs that exhibit unexpected complex patterns that are often missed by conventional detection tools, especially those involving nested or multi-breakpoint events. Experiments on simulated and real datasets demonstrate that gSV achieves competitive performance on canonical SV classes and improved sensitivity for complex SV patterns. Furthermore, applications of gSV to four breast cancer cell lines and a Chinese breast cancer cohort identified SVs in breast cancer-associated genes, further illustrating the potential utility of gSV.

In the following sections, we first detail the methodological framework of gSV and the experimental design for benchmarking in the Methods section. In the Results and Discussion section, we present a quantitative comparison of gSV with existing tools using simulated and real data. We further validate the unique findings of gSV against the existing literature, illustrating the usefulness of our new tool in genomic and clinical studies.

## Methods

### Detailed description of gSV

The detailed workflow of gSV is shown in Figure 1. We propose a novel *encoding* strategy that coverts information from reads/reference into a matrix form, as illustrated in Figure 1, which facilitates us to analyze one chromosome at a time. After encoding, the second step is *detecting* candidate regions with strong SV signals using the graph cut method [29]. In each candidate region, there may be multiple reads belonging to different subtypes, representing different SVs or genotypes, for which we carry out the *clustering*. Then we apply an *assembling* method (Wtdbg2) to generate a long consensus [23] that covers the synthesized information from multiple reads within each cluster. After zooming in from the whole genome to candidate regions, our assembling is computationally much cheaper. Moreover, with a longer consensus, gSV has a chance to compare the differences between the reference and the underlying sequence comprehensively. With only one consensus, we cannot afford missing detailed/complex signals by using global alignment tools such as minimap2 [14] and NGMLR [24]. Therefore, we adopt the idea of MEM [8] between the consensus and the reference in our *realigning* step. In the *finalizing* step, we transform realignment results into SV callings. By properly incorporating the assembling and the MEM method, gSV provides a step (that did not exist before) to revisit the missed or misspecified signals for correcting the final calling. In the following, we shall explain these steps in detail.

**Figure 1.**
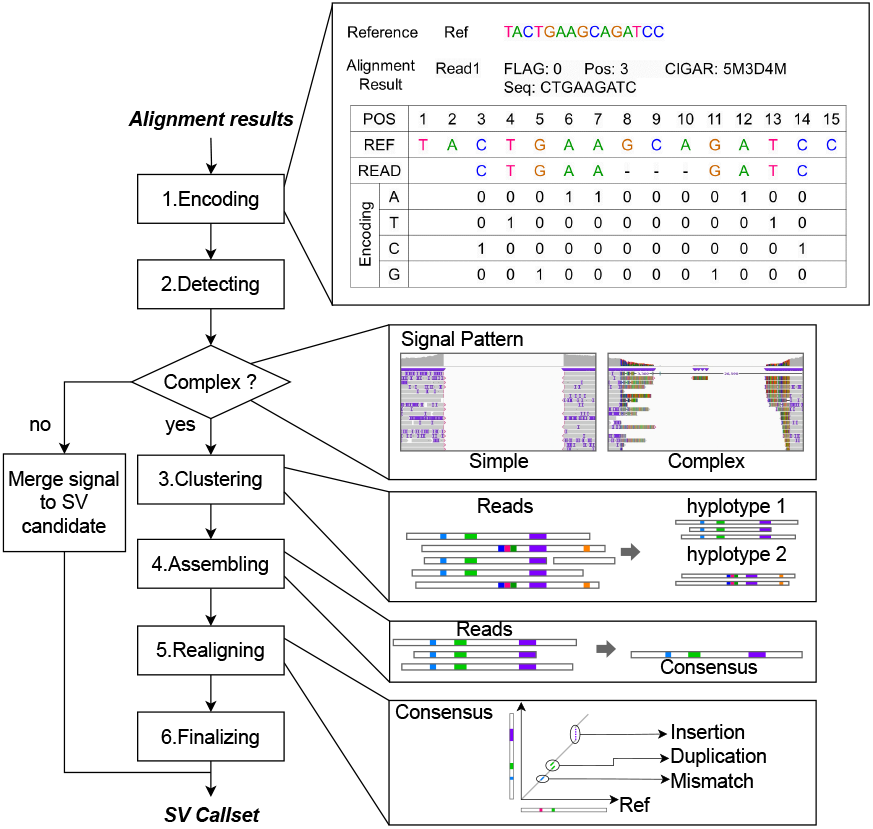
The workflow of gSV consists of six steps. Step 1. Encoding: converting reads/reference into a matrix representation to enable chromosome-scale analysis. Step 2. Detecting: using graph-cut method to detect candidate regions with strong SV signals. Step 3. Clustering: clustering reads within each complex candidate region into subtypes to resolve heterogeneous genotypes. Step 4. Assembling: generating long, accurate consensus sequences (via Wtdbg2) for each cluster. Step 5. Realigning: aligning the consensus to the reference using Maximal Exact Matches (MEMs) to capture discrepancies missed by global aligners. Step 6. Finalizing: transforming realignment results into SV callings.

#### Encoding

Unlike existing alignment-based SV detection tools that rely on predefined SV models, our method directly captures all differences between the reference genome and sequencing reads without pre-specifying SV types. These differences encompass sequencing errors and potential variant signatures. To enhance the ability to capture complex alignment patterns that are often lost in traditional CIGAR-based parsing, we adopt a matrix-based encoding method to capture base-level inconsistencies, positional gaps, and strand discordance in a unified numerical form. For each chromosome and read, we encode the reference and aligned read as 4 *× length* matrices representing nucleotides *A, T*, *C*, and *G*. Using the read’s reported genomic coordinates, we place the read matrix onto the reference coordinate system and compute their absolute difference. Because most bases match, the resulting matrices are sparse, enabling efficient memory usage and parallel processing.

Read matrices are generated from standard Sequence Alignment Map (SAM) fields (FLAG, POS, CIGAR, SEQ). Only essential alignment operations are retained in the main analysis: matches or mismatches (“M”/”=“/”X”), insertions (“I”), and deletions (“D”), while soft/hard-clipped bases are excluded. Supplementary alignments marking split reads are integrated with their primary alignments into a single matrix to preserve chimeric structure. Details of matrix construction, handling of CIGAR edge cases, and treatment of strand discordance are explained in Supplementary Note 1.3.

In addition, when encoding each read, every clear and easily recognizable signature will be recorded simultaneously. Taking the case of Step 1 in Figure 1 as an example, when encoding this read, we record a deletion with 3 bps in positions 8 to 10. The user can choose whether or not to use the prior information for subsequent SV detection. Different to other SV callers, our matrix-based encoding method preserves all inconsistencies, increasing the possibility of detecting SVs. To facilitate the subsequent discussion, let *S*_*j*_ =1 or 0 represents whether the *j*th position corresponding to the reference is a variant or not, *r*_*ij*_ = (*a*_*ij*_, *t*_*ij*_, *c*_*ij*_, *g*_*ij*_) indicates the encoding result for the *j*th position in the *i*th read, and *ref*_*j*_ indicate the encoding result for the *j*th position in the reference.

#### Detecting

After encoding the signals in a matrix form, we proceed to detect candidate regions with strong SV signals [29]. Using the above notations, we formulate the problem as follows:

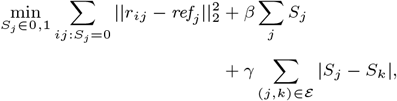

where ℰ indicates a neighboring set that satisfies |*j* − *k*| = 1.

The formula consists of three parts. The first part represents the loss when a position is labeled as non-variant (*S*_*j*_ = 0), and the *L*_2_ norm is used to quantify the difference between reads and the reference. The second part is a penalty when a position is labeled as variant (*S*_*j*_ = 1), with the parameter *β* acting as a threshold for supporting evidence. This model trades off between labels 1 and 0 based on the first two terms. To further smooth the label sequence and to encourage the detection of continuous variants, we incorporate a penalty for neighboring inconsistent labels at the third term. This formulation allows us to detect candidate regions with strong signals for subsequent analysis.

#### Clustering

In each candidate region, there may be multiple reads belonging to different subtypes, representing different SVs or genotypes within the region.

Users may optionally employ prior information derived from the encoding step to help in the detection. When this option is enabled (default setting), candidate regions are classified into two categories: regions with clear signatures and regions with complex or ambiguous signatures (usually containing complex SVs and some simple SVs with complex alignment patterns). If the option is disabled, all candidate regions are treated as regions with complex signatures, ensuring comprehensive detection of potential SVs.

For regions with clear signatures, prior information obtained from the encoding step usually contains only one or two consistent and unambiguous signatures. So we utilize frequency-based clustering to determine the subtype or genotype of SVs within these regions. Specifically, signatures with the same type, similar length and position are grouped together. By analyzing the number of supporting reads, we determine their genotypes and then make a final call of their corresponding SV.

For reads in regions with complex and ambiguous signatures, the alignment results are often too chaotic to discern what is happening in the regions. Therefore, we use assembly and alignment techniques to help figure out the sequence structure. But directly assembling reads corresponding to different subtypes together may lead to problematic consensus that does not reflect the features of any subtype. To solve this problem, we cluster the reads in such regions first.

Given that SVs usually cover many bases, if multiple subtypes of SVs exist, the reads corresponding to different subtypes exhibit significant deviations within the region. However, if only one subtype exists, the differences among reads are mainly due to other factors, such as sequencing or mapping errors. Importantly, we prefer only one subtype in one region, which is usually the case. This task differs somewhat from a classical clustering problem but fits within the framework of a statistical hypothesis test. The null hypothesis is that a pair of reads are from the same subtype, versus the alternative hypothesis that they are not from the same subtype.

To construct a test statistic, we directly calculate the differences between each pair of reads. The difference is based on the Levenshtein distance [19] within the candidate region. Specifically, we can apply a user-defined threshold to give the critical region. Then we select all the reads that appear in that region and arrange them in order of their lengths, from the longest read to the shortest read. We use the longest read as a representative sequence and test the similarity between each read and the representative. If the similarity score between a read and the representative sequence of a subtype falls below a user-defined threshold for the first time, we assume that the null hypothesis is invalid and that this read belongs to a different subtype. Then we create a new subtype and use this read as a representative sequence. Subsequent reads will be compared for similarity with representative sequences of different subtypes and assigned to the category with the highest similarity score. This algorithm runs efficiently with a time complexity of O(*n*), where *n* is the number of reads in the candidate region.

The Levenshtein distance, also known as the edit distance, is used to measure the difference between two sequences. Since sequences of different lengths will inevitably result in lower similarity scores, we use the left endpoint of the candidate region as the starting point to extract sequences of the same length for comparison. If there are reads that do not cover the left endpoint of the region, we will use the start position in the reference corresponding to the later read as the starting point to extract the sequence.

#### Assembling

For reads within a cluster, we utilize the assembly tool wtdbg2 [23] to generate a consensus. This assembling step enables us to obtain a longer contiguous sequence that covers the combined information from the reads within the cluster. Wtdbg2 is also used in DeBreak [3] to detect long insertions by assembling the reads around them. Please note that assembly is usually computationally expensive, which makes these assembly-type methods unpopular in SV detection. But assembly indeed provides an option to recover the sequence information for comprehensive checking, especially when the alignment information is complex. Thus how to economically use it is the key issue here.

Rather than using the assembly on raw reads, we apply Wtdbg2 only in the candidate region. In this way, we take advantage of the assembly-based methods and save unnecessary computation at the same time. Moreover, most SVs may be in a simple format, such as DELs, INSs, INVs, and DUPs. We also provide the option to skip the regions with simple SV signatures and only carry out the assembly on the candidate regions with complex SV signatures, which further reduces the computational burden.

#### Realigning

We align the assembled consensus back to the reference genome to determine the variants in the cluster. In this step, we note that SVDSS [6] also utilizes a similar procedure, but the purpose is to detect DELs and INSs due to the use of an improper alignment tool. It is important to mention that, we only have one consensus for each cluster. Common alignment tools like minimap2 [14] and NGMLR [24] aim to minimize the overall alignment loss by mapping the sequence to a specific part of the reference. Due to their mechanism, these tools may not achieve the minimum loss. In our case with only one consensus, we cannot afford such an error that may completely disrupt detection. Thus, we adopt the idea of MEM for the realigning step. MEM identifies and reports the maximum exact matches between the consensus and the reference while disregarding some gaps. This approach, despite yielding some gaps and multiple possible matches, keeps as much information as possible and allows us to infer the differences between the consensus and the reference more accurately. Moreover, MEM also enables us to capture information about potential complex SVs. In this paper, we utilize the copMEM2 [8] tool to carry out the MEM alignment.

#### Finalizing

In the finalizing step, the alignment results are transformed into the final SV callings. This step is similar to the encoding step, but with additional considerations for multiple possible matches and gaps therein.

When using MEM, both forward matches and inversion matches are reported, along with their positions in the consensus and reference sequences, as well as the match lengths. Due to various factors such as sequencing errors, mapping errors, SNPs, and small indels, gaps may be present in alignment results, apart from the SVs. These gaps are typically short, covering only a few bps. We will skip the gap not introduced by SVs, and convert inversion matches to the forward sequence. Subsequently, final SVs are determined based on the aforementioned encoding step. It is important to note that the final SVs may not necessarily fall into common SV types such as DELs and INSs. Therefore, we infer complex SV types directly from the MEM results.

### Performance evaluation

We use both simulated and real datasets to comprehensively assess SV detection performance. We compare gSV with five state-of-the-art SV callers: PBSV [21], Sniffles [26], cuteSV [11], DeBreak [3], and SVision-pro [28]. First, we conduct benchmark comparisons using simulated data, which provides controlled scenarios to quantify recall and precision across different SV types and complexities. Next, we perform benchmark comparisons using a real HG002 dataset, whose ground truth includes high-confidence INSs and DELs. To extend the evaluation across diverse SVs and real-world genomic complexity, we further test our method on additional datasets, including family trios for Mendelian inheritance validation[5], whole-genome sequencing of breast cancer cell lines[12], and population-scale breast cancer cohorts for clinical utility analysis. However, as these data lack a comprehensive ground truth, we perform manual inspection using the Integrative Genomics Viewer (IGV) to confirm alignment patterns, breakpoint support, and concordance with other evidence (e.g., read-depth signals).

#### Simple SV and Complex SV benchmark

We use simulated data and HG002 as benchmarks for simple SVs to evaluate the performance of different methods. For simulated data, we utilize the simulator VarSim [18] to randomly insert variants from the Database of Genomic Variants (DGV) (2,675 INSs, 4,972 DELs, 1,409 DUPs, and 362 INVs) into the reference genome (GRCh37). Then we utilize PBSIM2 [20] to simulate Continuous Long Reads (CLRs). For HG002, we use the BAM file obtained by aligning SMRT sequencing data to the hs37d5 reference using pbmm2 (the minimap2 wrapper), and we utilize the Tier 1 benchmark set (including highly confident INSs and DELs) as the ground truth. The size distributions of SVs in simulated data and HG002 are as shown in Supplementary Figure 1.

We use simulated complex SVs as the benchmark to evaluate the performance of different methods. Based on previous studies on complex SVs [4; 17], we classified complex SVs into five major categories with 12 subclasses according to their structural composition and simulated them (shown in Fig. 2). The first three combinations (ID 1-3) belong to one category, formed by linking adjacent simple subsidiary SVs together. The last two combinations (ID 4-5) belong to the second category, composed of overlapping and nested sub-SVs. Following the idea of creating complex SVs used in Sniffles and SVision, we utilize VISOR [2] to construct complex SVs and PBSIM2 to simulate reads.

**Figure 2.**
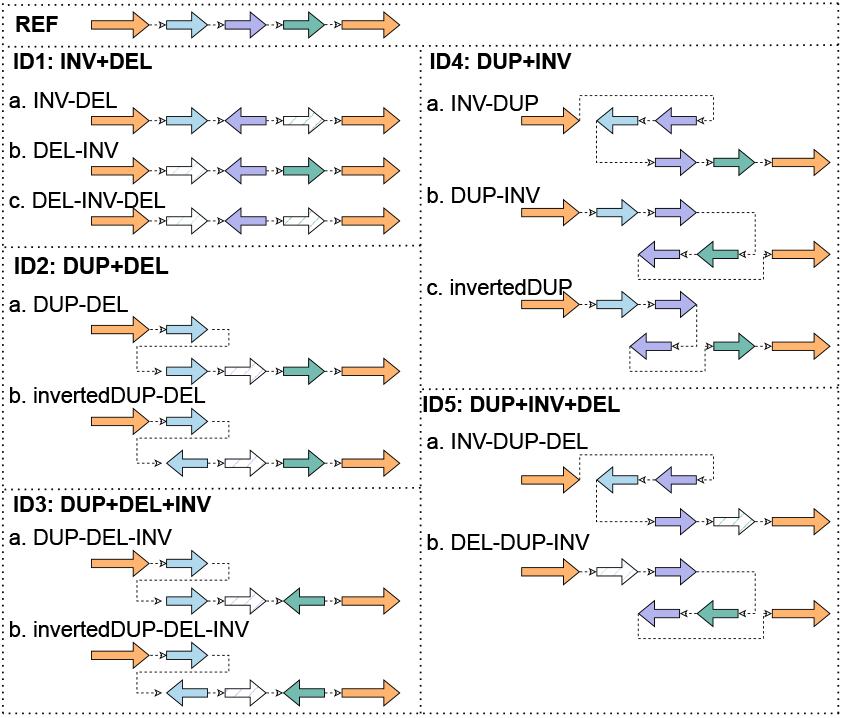
Diagrams of simulated complex SVs. IDs 1–3 are constructed by linking adjacent simple sub-SVs. IDs 4–5 are composed of overlapping or nested sub-SVs.

We use Truvari [7] to evaluate the results of gSV, PBSV [21], Sniffles [24], cuteSV [11], DeBreak [3] and SVision-pro [28] on simple and complex SVs. For complex SVs, we evaluate the different tools from both the regional and type perspectives. We first focus on evaluating whether the tools can identify the regions correctly. Since SVision-pro is the only tool capable of reporting complex SVs, we compare our gSV exclusively with it. We extract the start and end positions of each complex SV and use Truvari to calculate recall, precision, and F1 scores. Furthermore, we evaluate the detection results from the perspective of SV sub-types. Since all tools, other than SVision-pro, can only detect the parts of complex SVs, we divide each complex SV into its constituent simple sub-type SVs, which form a new ground truth dataset. In other words, we examine each sub-component of complex SVs for position, size, and type. For PBSV, Sniffles, cuteSV, and DeBreak, we directly compare the detection results with the split ground truth set. For gSV and SVision-pro, we split the detected complex SVs and merge them with the detected simple SVs to perform the evaluation.

#### Mendelian consistency analysis

For real data lacking ground truth, traditional metrics (e.g., recall, precision, and F1-score) cannot be directly applied to evaluate the detection results. Here we leveraged Mendelian consistency in family trios as a surrogate validation framework, where fewer observed Mendelian inheritance violations directly correlate with reduced false-positive rates. Mendelian consistency is a statistical measure used to evaluate whether observed SVs in a family trio (e.g., parents and offspring) adhere to Mendelian inheritance laws. It is typically calculated as the proportion of SVs in offspring that are genetically compatible with parental genotypes: (Total SVs - Mendelian violations SVs)/ Total SVs.

We perform Mendelian consistency analysis on HiFi sequencing data from four previously published families: the Chinese Trio, the Yoruba Trio, the Southern Han Chinese Trio, and the Puerto Rican Trio. We use Minimap2 to align the sequencing reads to the GRCh38 reference genome. Five variant callers are employed: gSV, cuteSV, DeBreak, SVision-pro, and Sniffles. For the three callers without built-in merging capabilities (gSV, cuteSV, and DeBreak), we first performed independent variant calling for each sample, and then compared the results of two merging methods (Jasmine[13] and SURVIVOR[10]). To eliminate the impact of de novo SVs, we retained only SVs detected in at least two samples within each trio (e.g., parent-parent or parent-child pairs). Finally, we used the Mendel plugin in BCFtools to assess Mendelian consistency.

#### Tumor-normal paired cell line SV analysis

To explore novel SV loci and complex structures, we examine four open-source breast cancer tumor-normal paired cell line datasets: HCC1395, HCC1937, HCC1954, and HG008. Sequencing reads are aligned to the GRCh38 reference sequence using minimap2. And we use gSV, cuteSV, DeBreak, SVision-pro, and Sniffles for SV detection.

Due to the lack of ground truth, we performed manual inspection using IGV to visually inspect our unique findings. Given the impracticality of manually vinspecting more than 20,000 SVs typically detected per sample, we implement a pipeline to prioritize candidate SVs. First, we analyze SV callsets across multiple tools using Jasmine to compare the overlap of detected SVs and identify the subset of tools supporting each SV. Next, we annotate all SVs with ANNOVAR[27] to prioritize functionally relevant variants (e.g., genes with potential clinical implications, exonic regions). Finally, we focus validation efforts on SVs uniquely identified by gSV to assess its ability to detect novel or challenging SVs missed by existing methods.

#### In-house breast cancer data analysis

We further analyze an in-house long-read sequencing dataset acquired at the People’s Hospital at Peking University from 234 samples (188 samples are from breast cancer patients, and the other 46 are from people currently not diagnosed with breast cancer). Focusing on breast cancer, we develop a 28-related-gene (more details see Supplementary Table 1) panel for Pacific Biosciences platform-based sequencing. We apply gSV together with PBSV, Sniffles, cuteSV, DeBreak, and SVision-pro to detect SVs from each sample. We first filter out the detected SVs shorter than 50 bps. For comparison, an important step is to determine the intersections of the SVs detected by the tools. Given that the SVs detected by different tools may be slightly different in terms of position and/or length, we give some tolerance as follows:

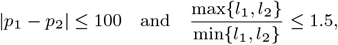

where *p*_1_ and *p*_2_ represent the start positions for the detected SVs and *l*_1_ and *l*_2_ represent their lengths. For each type of SV, we go through each pair of candidate SVs from the different tools and determine whether they are identical. Then, across samples, we follow a similar procedure to determine whether an identical SV exists in different samples. After that, we performed manual inspection using IGV to confirm our unique findings.

## Results and Discussions

In this section, we present comparative results in different data sets and discuss the possible biological and clinical significance of the unique findings of gSV in breast cancer data.

### Performance evaluation of simple and complex SVs in datasets with known ground truth

For simple SVs, as shown in panel (a) of Fig. 3, gSV outperforms the other tools for all read depths in terms of recall, precision, and F1-score. Panel (b) of Fig. 3 compares the performance of different tools across different types of simple SVs. For INSs, the alignment pattern is relatively simple, so most tools show comparable performance. For DELs, gSV achieves a moderate improvement over state-of-the-art tools, with an approximate 0.7% increase in F1-score compared to the best-performing existing method. In contrast, this advantage becomes markedly more pronounced for DUP and INV, where gSV demonstrates F1-score improvements of 3.1% and 4.0%, respectively. This is because the alignment patterns of these SV types exhibit more complex scenarios, as illustrated in panels (d) and (e) in Figure 3. Specifically, panel (d) shows a DEL where the corresponding region does not exhibit a clean deletion pattern but displays a discontinuous alignment with numerous mismatches. Panel (e) shows a large-scale DUP where the variant signal is not directly discernible in most reads. Only a few of split-read alignments exhibit partial truncation on the right flank, indicative of the DUP breakpoint. However, there is a pronounced elevation in read depth across the duplicated region. These patterns are different from those expected for simple DELs or DUP in alignment models, rendering them undetectable by existing tools. This further demonstrates the unique capability of our method to resolve SVs with complex alignment patterns. In addition, we apply gSV to the HG002 dataset (only containing INSs and DELs) from the Genome in a Bottle (GIAB) consortium [31], and find that gSV performs similarly well to the benchmarking tools for INSs and DELs (panel (c) of Fig. 3), consistent with the results shown in panel (b).

**Figure 3.**
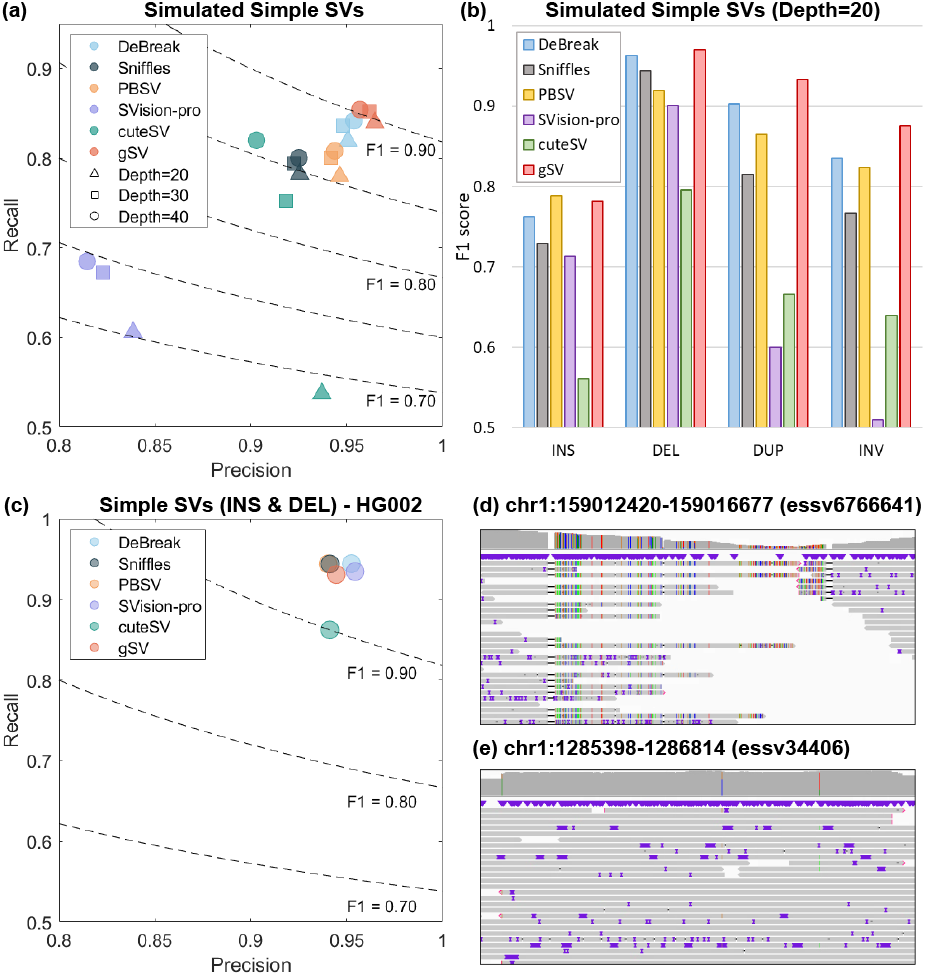
Panel (a) shows the performance of different methods in detecting simple SVs in terms of recall, precision, and F1-score. gSV uniformly performs better in all depths. Panel (b) compares the F1-scores of different methods for detecting specific types of simple SVs. gSV demonstrates a more pronounced advantage in detecting DUP and INV compared to existing methods. Panel (e) evaluates the performance of different methods in detecting simple SVs (INS and DEL) on the HG002 benchmark dataset. gSV performs similarly well to the benchmarking tools in INSs and DELs, consistent with the results shown in Panel (b). Panels (d) and (e) show the IGV screenshot of a DEL and a DUP uniquely detected by gSV. Both variants exhibit complex alignment patterns, which prevented their detection by other tools. (The contents in parentheses indicate their identifiers in the DGV.)

For complex SVs, the left panel of Figure 4 shows that gSV achieves substantially higher accuracy than existing tools in both breakpoint localization and SV sub-type determination. More results for other types of simulated complex SVs are shown in Supplementary Figure 2. The right panel shows the numbers of complex SVs detected by SVision-pro and gSV across different types, with red representing gSV and blue representing SVision-pro. For the INV+DEL(ID1) and DUP+DEL+INV(ID3) categories, the numbers of CSVs detected by SVision-pro and gSV differr(striped bars), but only to a limited extent. In contrast, for the remaining three categories, particularly the highly complex DUP+INV(ID4) and DUP+INV+DEL(ID5) types that involve overlapping or nested sub-SVs, SVision-pro detects markedly fewer CSVs than gSV. Among the detected CSVs, SVision-pro and gSV show comparable positional precision, as shown in Supplementary Figure 2. However, in absolute terms, gSV identifies more true CSV loci. When the accuracy of CSV type is taken into consideration, gSV exhibits a clear advantage.

**Figure 4.**
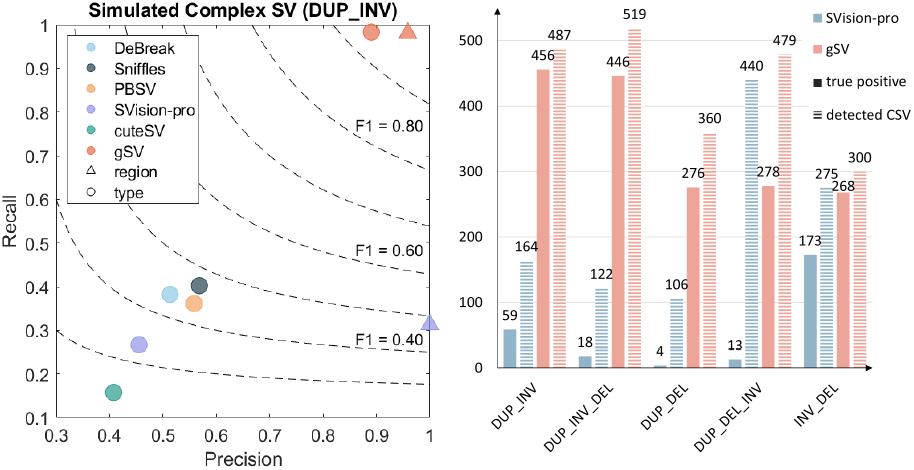
The left panel shows the performance of different methods in detecting one example type (ID4: DUP-INV) of complex SV by evaluating the region and type, respectively. gSV demonstrates better performance resolving these complex cases, underscoring its capability to accurately detect and characterize multiple breakpoints or nested structural variants, which are often missed by conventional detection tools. The right panel shows the numbers of complex SVs detected by SVision-pro and gSV across different types (striped blue for SVision-pro and striped red for gSV), and the corresponding numbers of true positive SVs (solid blue for SVision-pro and solid red for gSV).

### Mendelian consistency assessment in family trios

Figure 5 shows the Mendelian consistency of different tools in four family trio data. We can see that gSV demonstrates greater Mendelian consistency compared to other tools, reflecting its higher precision in SV detection.

**Figure 5.**
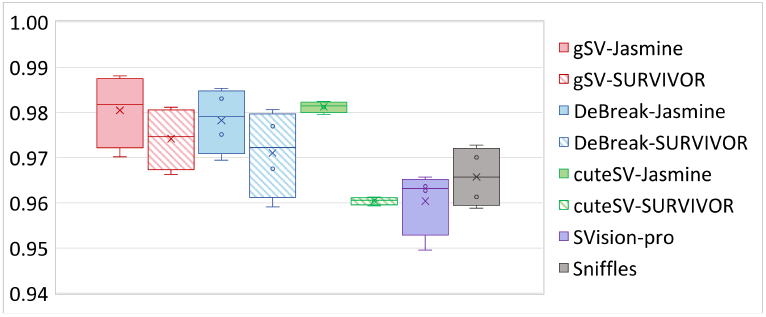
Mendelian consistency of different tools in four family trio data. gSV demonstrates higher Mendelian consistency compared to other tools.

### SV analysis in breast cancer tumor-normal cell lines

Following the analytical framework in Section 2.2, we obtain the gSV-unique-detected SVs and then perform literature validation to corroborate their biological relevance. Panel (a) of Figure 6 demonstrates a somatic DEL in HCC1954-tumor only detected by gSV, located within the 5-Hydroxytryptamine Receptor 1A (HTR1A) exonic region. Published studies [16] indicate that HTR1A significantly inhibits triple-negative breast cancer (TNBC) cell development in vivo and in vitro through downregulation of both canonical and noncanonical TGF-*β* pathways. Notably, the expression level of HTR1A was significantly lower in TNBC tissues compared with that in paracancerous tissues, and knockdown of HTR1A significantly enhanced the migration and invasion of MDA-MB-231 and Hs578T breast cancer cell lines. Panel (b) of Figure 6 demonstrates a germline DUP in the HCC1937 cell line exclusively identified by gSV. This SV resides within the FLG exonic region, which encodes filaggrin. The partial exonic DUP disrupts the filaggrin architecture, potentially leading to skin barrier function, which may lead to higher cancer susceptibility [25]. Another studies show that FLG were found amplified in 12.7% of breast tumors [1]. Additional gSV-unique-detected SVs localized to exonic regions of breast cancer-associated genes are provided in Supplementary Figures 4-11. Furthermore, Supplementary Figures 12-14 show exon-proximal upstream SVs in breast cancer-associated genes. Although these non-coding alterations do not directly disrupt core coding sequences or protein domains, they may alter the binding sites of transcription factors. Such regulatory perturbations could drive oncogenic progression through dynamic upregulation or downregulation of critical gene expression programs. Supplementary Figures 15-17 show gSV-unique-detected SVs in exonic regions of genes not currently linked to breast cancer. These findings provide potential candidates for investigating variations in cancer genomics, even in the absence of established oncogenic associations.

**Figure 6.**
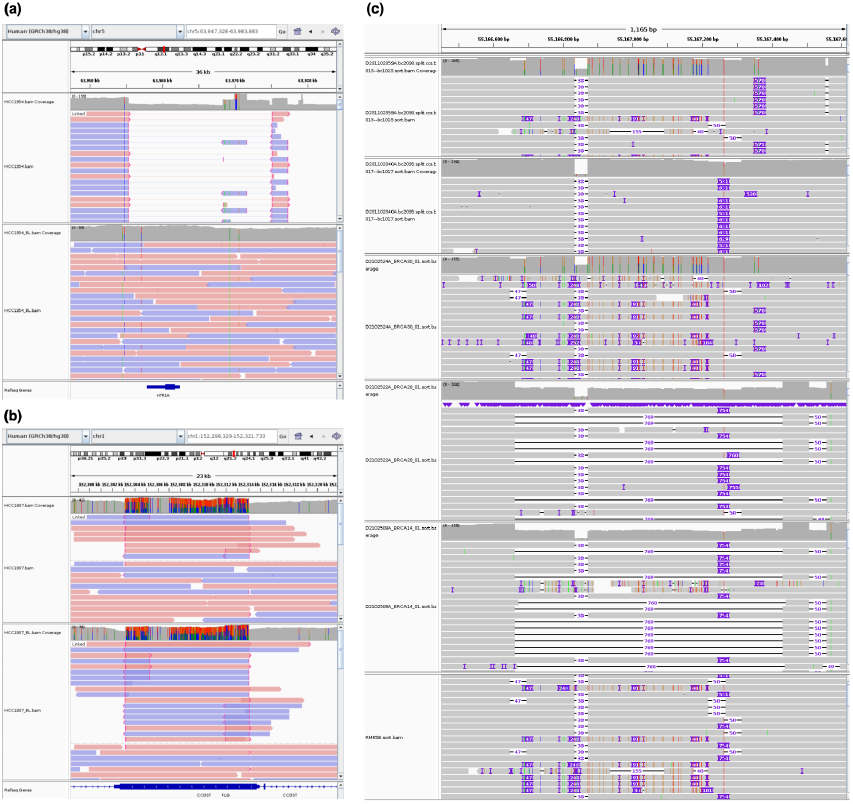
Panel (a) and (b) are the IGV screenshots of the SVs only detected by our gSV in HCC1954 and HCC1937. Panel (c) shows a DUP (Chr7:Duplication at EGFR in chromosome 7: 55167022-55167776) uniquely detected by gSV in six breast cancer patients, which shows an American College of Medical Genetics and Genomics (ACMG) class of 3 (unclear breast cancer risk).

### SV analysis of in-house breast cancer data

In this in-house dataset, we analyzed targeted sequencing data of 28 breast cancer–related genes from 234 samples, including 188 breast cancer patients and 46 individuals without a breast cancer diagnosis. Under this setting, gSV detects 8 unique SVs that are not detected by the other tools. We show one such example of a DUP in EGFR in panel (c) of Figure 6. It shows an American College of Medical Genetics and Genomics (ACMG) class of 3, which means an unclear risk related to breast cancer [22]. The same SV is detected in five other patients but not in healthy people, indicating a possible association with breast cancer. Details of the remaining gSV-unique-detected SVs are provided in Supplementary Table 2. Although these variants are located in non-exonic regions, their translational implications warrant further investigation, as they may reside in critical regulatory elements influencing gene expression, splicing mechanisms, or epigenetic modifications that could underlie phenotypic variability or disease susceptibility.

## Conclusion

Our proposed gSV inherits high sensitivity from alignment-based methods and high specificity from assembly-based methods. The inclusion of MEM further improves its capability in capturing detailed and complex signals. This hybrid design enables gSV to detect both simple SVs and complex SVs formed by non-prespecified combinations of basic SV types. Benchmark evaluations on simulated data with ground truth show its superior performance in recall and precision compared to other existing popular tools. When we apply the new tool to detect SVs in public datasets and in-house breast cancer dataset, we obtain promising findings not detected by existing tools. Overall, gSV provides a general and easy-to-interpret pipeline for detecting SVs.

## Declaration

### Data availability Statement

Sample HG002, HG008, HCC1395, HCC1937, HCC1954, have been consented for commercial use and for public posting of Personally Identifying Genetic Information (PIGI), which allows open-access, public posting of extensive genetic data. The studies involving human participants related to breast cancer data were reviewed and approved by Peking University People’s Hospital ethics committee. The patients/participants provided their written informed consent to participate in this study.

### Availability of data and materials

HG002 data was downloaded from https://ftp-trace.ncbi.nlm.nih.gov/giab/ftp/data/AshkenazimTrio/HG002_NA24385_son/PacBio_CCS_15kb/alignment/. The Tier1 benchmark SV callset and high-confidence HG002 region were obtained from https://ftp-trace.ncbi.nlm.nih.gov/giab/ftp/data/AshkenazimTrio/analysis/NIST_SVs_Integration_v0.6/. HG008 normal-tumor paired cell data were downloaded from https://ftp-trace.ncbi.nlm.nih.gov/ReferenceSamples/giab/data_somatic/HG008/Liss_lab/PacBio_Revio_20240125/. HCC1395 normal-tumor paired cell data were downloaded from https://downloads.pacbcloud.com/public/revio/2023Q2/HCC1395/. HCC1937 normal-tumor paired cell data were downloaded from https://trace.ncbi.nlm.nih.gov/Traces/?view=run_browser&acc=SRR28305185&display=metadata and https://trace.ncbi.nlm.nih.gov/Traces/?view=run_browser&acc=SRR28305182&display=metadata. HCC1954 normal-tumor paired cell data were downloaded from https://trace.ncbi.nlm.nih.gov/Traces/?view=run_browser&acc=SRR28305163&display=metadata and https://trace.ncbi.nlm.nih.gov/Traces/?view=run_browser&acc=SRR28305160&display=metadata. The human reference genome GRCh37 was downloaded from http://ftp-trace.ncbi.nih.gov/1000genomes/ftp/technical/reference/phase2_reference_assembly_sequence/. The human reference genome GRCh38 was downloaded from https://ftp-trace.ncbi.nlm.nih.gov/ReferenceSamples/giab/release/references/GRCh38/. Sequencing data for in-house breast cancer samples have been deposited in the Genome Sequence Archive for Human (GSA-Human) at the National Genomics Data Center (NGDC) under accession codes HRA003120, HRA003172, and HRA008465.

### Code availability

gSV is publicly available at https://bioinformatics.hkust.edu.hk/Software.html or https://github.com/jhaoae/gSV.

### Author contributions statement

J. Hao and J. Shi developed and implemented the analysis algorithms, carried out data analysis, and drafted the paper. S. Lian, Z. Zhang, and Y. Luo helped improve the analysis methods. S. Wang, X. Fan, T. Ishibashi, and W. Yu conceived the idea, designed and supervised the study. T. Hu and D. Wang coordinated sample collection and data acquistion. All authors helped to proofread and revise the manuscript.

**Jingyu Hao** is a PhD candidate in the Department of Electronic and Computer Engineering at the Hong Kong University of Science and Technology. Her research interests lie in third-generation sequencing data analysis and structural variant detection.

**Jiangdong Shi**, PhD, is a Postdoctoral Fellow in the Department of Statistics and Data Science, The Chinese University of Hong Kong. His research interests include Meta-analysis, Bayesian analysis, and third-generation sequencing data analysis.

**Sheng Lian** received his PhD degree in Statistics from The Chinese University of Hong Kong. He is the co-founder of SinoReco Technology.

**Zhen Zhang**, received his PhD degree in Statistics from The Chinese University of Hong Kong. He is currently affiliated with the Kunming Institute of Physics.

**Yongyi Luo** is a PhD candidate in the Department of Statistics and Data Science, The Chinese University of Hong Kong. Her research interests lie in statistical inference, sequencing data analysis, and healthcare-related topics.

**Taobo Hu**, PhD (Life Sciences), MD, is currently affiliated with Peking University People’s Hospital.

**Toyotaka Ishibashi**, PhD, is an Associate Professor in the Division of Life Science at the Hong Kong University of Science and Technology. He is interested in Biophysics, Chromatin structure, and Transcription dynamics.

**Depeng Wang** is Chief Executive Officer (CEO) of GrandOmics. He has been engaged in research on Chinese structural variation data and participated in several major research projects based on third-generation sequencing data.

**Shu Wang**, PhD, is the director of the Breast Center at Peking University People’s Hospital, where she also serves as a professor and doctoral supervisor.

**Xiaodan Fan**, PhD, is a professor of Statistics, The Chinese University of Hong Kong. He is interested in probabilistic modeling, statistical computing, pattern recognition, and computational biology/ bioinformatics.

**Weichuan Yu**, PhD, is a professor in the Department of Electronic and Computer Engineering at the Hong Kong University of Science and Technology. He is interested in computational analysis problems with biological and medical applications.

