## Supplementary Notes, Tables, Figures for "gSV: a general structural variant detector using the third-generation sequencing data"

### Supplementary Materials:
### *g*SV: a *g*eneral structural variant detector using the third-generation sequencing data


Jingyu Hao[a,1], Jiandong Shi[b,1], Sheng Lian[a], Zhen Zhang[a], Yongyi Luo[b], Taobo Hu[c], Toyotaka Ishibashi[d], Depeng Wang[e], Shu Wang[c,*], Xiaodan Fan[b,*] and Weichuan Yu[a,*]

[a]*Department of Electronic and Computer Engineering, Hong Kong University of Science and Technology, Hong Kong SAR, China*
[b]*Department of Statistics, The Chinese University of Hong Kong, Hong Kong SAR, China*
[c]*Department of Breast Surgery, Peking University People's Hospital, Beijing, China*
[d]*Division of Life Science, Hong Kong University of Science and Technology, Hong Kong SAR, China*
[e]*GrandOmics Inc, Beijing, China*


#### 1. Supplementary Notes

##### 1.1. Data simulation

***Simple SV simulation***

To simulate simple structural variations (SVs) more realistically, we utilized the simulator VarSim[1], which can sample from an existing database, to insert four types of simple variations (deletion (DEL), duplication (DUP), insertion (INS), and invertion (INV)) into the reference genome (GRCh37). These variations were sourced from the Database of Genomic Variants (DGV), which summarizes the structural variations in the human genome. We construct the SV set according to the proportion of different SVs in DGV. Since VarSim does not support translocation (TRA), and INV appears relatively rarely in the DGV database, we increased the proportion of INV to 4% to better evaluate the detection capabilities of different callers for INV.

Specifically, we generate 2,700 INSs, 5,700 DELs, 1,530 DUPs, and 400 INVs. However, practical issues such as sampling multiple SVs at the same position limited the generation of some SVs, resulting in the following numbers of simple SVs generated: 2,675 INS, 4,972 DELs, 1,409 DUPs, and 362 INVs. The commands to generate the simulation data are shown as follows:

- `wget http://web.stanford.edu/group/wonglab/varsim/insert_seq.txt`
- `wget http://web.stanford.edu/group/wonglab/varsim/GRCh37_hg19_supportingvariants_2013-07-23.txt`
- `varsim.py --vc_in_vcf vc_file --sv_insert_seq insert_seq.txt --sv_dgv GRCh37_hg19_supportingvariants_2013-07-23.txt --reference hs37d5.fa --id sim --seed 17 --disable_sim --vc_num_snp 0 --vc_num_ins 0 --vc_num_del 0 --vc_num_mnp 0 --vc_num_complex 0 --sv_num_ins 2700 --sv_num_del 5700 --sv_num_dup 1530 --sv_num_inv 400 --sv_percent_novel 0 --sv_min_length_lim 50 --sv_max_length_lim 10000 --nlanes 1 --total_coverage 1 --java_max_mem 50g --simulator_executable $OPT_DIR/ART/art_bin_VanillaIceCream/art_illumina --out_dir out --log_dir log --work_dir work`

***Complex SV simulation***

We utilized VISOR [2] to construct complex SVs. Firstly, we used the built-in script of VISOR to randomly generate some basic SVs. Taking ID 5 (DUP+INV+DEL) as an example, we first randomly generated duplications.

- `Rscript randomregion.r -d chrom.dim.tsv -n num -l length_mean -s length_sd -x exclude_regions.bed -v 'tandemduplication' -r '100' | sortBed > DUP.bed`

Next, based on the position of each DUP, we manually generated the Browser Extensible Data (BED file) containing the positions of INVs and DELs ('INV_DEL.bed'). Since VISOR does not support repeating insertions of SVs at

*Corresponding author(s).
[1]These authors contributed equally to this work.

overlapping positions, we first utilized ‘DUP.bed’ to insert duplications into the reference sequence, resulting in Sequence 1. Then, we insert INVs and DUPs into Sequence 1 based on ‘INV_DEL.bed’ and obtained the final sequence.

- `VISOR HACK -g ref.fa -bed DUP.bed -o /path/to/output`
- `VISOR HACK -g s1.fa -bed INV_DEL.bed -o /path/to/output`

It is notable that for the first three combinations (ID 1-3) of complex SVs (no overlapping relationship between sub-SVs), the sequence containing all SVs can be generated at once.

***Reads generation***

After obtaining the sequences with inserted SVs, we utilized PBSIM2 [3] to simulate Continuous Long Reads (CLRs). The quality scores of reads are generated by the factorized information criteria(FIC)-HMM of P6C4 chemistry.

- `pbsim  -hmm_model data/P6C4.model   sequence.fa`

#### 1.2. Software Version and Parameters

**Truvari (V4.1.0) [4]**

- `bgzip caller.vcf && tabix caller.vcf.gz`
- `truvari bench -b ground_truth.vcf.gz -c caller.vcf.gz -o dir --passonly -p 0.00 -r 1000 -P 0.5`

**Sniffles (V2.2) [5]**

- `sniffles -i sample.sorted.bam -v sniffles.vcf -reference reference.fa`

**CuteSV (V2.0.3) [6]**

- `cuteSV sample.sorted.bam reference.fa cutesv.vcf workdir`

**PBSV (V2.9.0) (https://github.com/PacificBiosciences/pbsv/)**

- `pbsv discover sample.sorted.bam sample.svsig.gz && pbsv call reference.fa sample.svsig.gzpbsv.vcf`

**Debreak (V1.0.2) [7]**

- `debreak -bam sample.sorted.bam -outpath debreak -rescue_large_ins -rescue_dup -poa -ref reference.fa`

**SVision-Pro (V1.8) [8]**

- `SVision-pro --target_path sample.sorted.bam --genome_path reference.fa --model_path {SVISIONPRO_DIR}/src/pre_process/model_liteunet_256_8_16_32_32_32.pth --out_path workdir --sample_name svision-pro --detect_mode germline`

**Jasmine (V1.1.5) [9]**

- `jasmine file_list=sample_list.txt out_file=Jasmine.merge.vcf`

**SURVIVOR (V1.0.7) [10]**

- `SURVIVOR sample_list.txt 1000 1 1 1 0 50 SURVIVOR.merge.vcf`

**Table 1**
Common FLAG meanings in alignment results (SAM file).

| Value | Meaning |
|---|---|
| 0 | Single-end sequencing, successfully mapped, and aligned to the forward strand. |
| 16 | Reverse-complemented relative to the reference genome (aligned to the reverse strand). |
| 2048 | Supplementary alignment: part of a chimeric alignment (e.g., split reads spanning distant genomic regions). |
| 2064 | Both a supplementary alignment and reverse-complemented. |

#### 1.3. Encoding Step in *g*SV

The encoding step aims to identify all differences between the reference genome and aligned reads, encompassing both sequencing errors and potential variant signatures.

Unlike existing alignment-based SV detection tools, we do not initially specify SV types and build corresponding models. Instead, we identify all the differences between the reference genome and the reads from the beginning. The reference sequence for each chromosome is encoded as a $4 \times reference\_length$ matrix, and the alignment results of each read are encoded as a $4 \times read\_length$ matrix. The start and end positions of each result are also recorded to determine their positions in the reference. By subtracting the two matrices, we obtain all the absolute differences between the reads and the reference. Unlike other approaches that capture signals conforming to predefined models via CIGAR strings, this matrix-based encoding preserves all potential variant information, making it possible to detect novel SVs whose patterns have not yet been summarized in existing models. Moreover, since most bases between the reads and the reference are identical, the resulting matrices are inherently sparse. This sparsity substantially reduces memory usage and computational cost, while enabling efficient parallel processing.

Specifically, the read matrices are constructed by extracting key fields from alignment results stored in the Sequence Alignment/Map (SAM) format, including the FLAG (a bitwise integer encoding alignment properties such as paired-end status and strand orientation), POS (the 1-based leftmost genomic position of the alignment), CIGAR string (a Compact Idiosyncratic Gapped Alignment Report summarizing alignment operations), and SEQ (the nucleotide sequence of the read). The CIGAR string comprises integer-operation pairs where the integer specifies the number of bases, and the operation defines the alignment type: "M" (generic alignment match, encompassing "=" for exact matches and "X" for mismatches), "I" (insertion relative to the reference), "D" (deletion relative to the reference), "S" (soft-clipped unaligned bases), and "H" (hard-clipped unaligned bases). Common FLAG values and their interpretations are detailed in Table 1.

Each sequencing read is encoded into a matrix with four rows corresponding to nucleotides $A$, $T$, $C$, and $G$. For the simplest case (FLAG = 0), the sequence is extracted from the SEQ field and encoded according to its CIGAR string. For CIGAR operations "M" (including "=" and "X"), each base is represented as a one-hot vector (e.g., $A \rightarrow [1\ 0\ 0\ 0]^T$). For CIGAR "I", the inserted bases are compressed into one column and added to the vector at the insertion position. For example, an 8-base insertion ("8I") $ATCGATCG$ at a $C$ nucleotide position would generate a vector $[2\ 2\ 3\ 2]^T$, calculated as $[2\ 2\ 2\ 2]^T$ (insertion) $+[0\ 0\ 1\ 0]^T$ (insertion position base $C$). For CIGAR operation "D", the corresponding column is encoded as a zero vector $[0\ 0\ 0\ 0]^T$. Bases corresponding to "S" or "H" are skipped because they are not part of the alignment. For FLAG = 16 (reverse-complement alignment), the SEQ field shows the reverse complement of the original read, but the same encoding approach is employed without additional transformations. Supplementary alignments (marked by FLAG=2048 or 2064) represent chimeric alignment records for a single read that cannot be fully represented by a single linear alignment. These occur when a read is split into multiple segments, resulting in one primary alignment (the highest-confidence alignment) and one or more supplementary alignments, each mapped to distinct regions of the reference genome. To preserve the chimeric relationship inherent in split-read alignments, we encode both the primary alignment and its supplementary counterparts into a single matrix. When supplementary and primary alignments share the same strand orientation, they are placed in the matrix based on their relative position, and the regions between them are represented by 0. For supplementary alignments on the opposing strand relative to the primary alignment, the corresponding segment in the supplementary SEQ is reverse-complemented before encoding to retain strand discordance as a potential variant signal.

##### *Toy Example Demonstration*

This section demonstrates how *g*SV's encoding operates through four toy reads representing the four alignment scenarios in the main text.

Suppose that we have the reference genome sequence $AAGCTTCTCACCCTG$ and four toy reads as follows:

- read 1: $TTCATCGATCGTCCCTG$ (aligned as given)
- read 2: $CAGGGGAATT$ (aligned in reverse complement orientation)
- read 3: $AAGCCCCTG$ (split-aligned)
- read 4: $AAGCTTCTCCAGGGT$ (chimeric aligned)

Figure 1 shows the alignment results where lowercase bases denote clipped regions. The corresponding Sequence Alignment/Map (SAM) format is shown in Figure 2, which is the standard format to record alignment results. Note that BAM files (Binary Alignment/Map), the compressed binary representation of SAM files, serve as the standard input format for our $g$SV pipeline.

```
Coor     1  2  3  4  5  6  7                           8  9  10 11 12 13 14 15
ref      A  A  G  C  T  T  C  *  *  *  *  *  *  *  *  T  C  A  C  C  C  T  G

+ read1              T  T  C  A  T  C  G  A  T  C  G  T  C  *  *  C  C  T  G
- read2  A  A  *  *  T  T  C                           *  *  *  C  C  C  T  G
+ read3  A  A  G  C  .  .  .                           .  .  .  C  C  C  T  G
+ read4  A  A  G  C  T  T  C                           T  C  c  a  g  g  t
- read4                                                      A  C  C  C  T  G  g  a  g  a  a  g  c  t  t
```

Figure 1: Alignment results of four reads. '+' denotes forward strand alignment; '-' denotes reverse complementary strand alignment.

| QNAME | FLAG | RNAME | POS | CIGAR | SEQ |
|---|---|---|---|---|---|
| read1 | 0 | ref | 5 | 3M8I2M2D4M | TTCATCGATCGTCCCTG |
| read2 | 16 | ref | 1 | 2M2D3M3D5M | AATTCCCCTG |
| read3 | 0 | ref | 1 | 4M5S | AAGCCCCTG |
| read3 | 2048 | ref | 11 | 4S5M | AAGCCCCTG |
| read4 | 0 | ref | 1 | 9M6S | AAGCTTCTCCAGGGT |
| read4 | 2064 | ref | 10 | 6M9S | ACCCTGGAGAAGCTT |

Figure 2: The alignment results of four reads in SAM format. The QNAME (query name) serves as the identifier for each read; the FLAG indicates alignment information (0: a primary alignment in the forward strand, 16: a reverse complement alignment, 2048: a supplementary alignment on the forward strand, and 2064: a supplementary alignment on the reverse strand); the RNAME (reference name) specifies the reference sequence to which the read is aligned; the POS indicates the starting position of the alignment on the reference, the CIGAR string describes the alignment specifics (M: matches or mismatch, I: insertions, D: deletions, and S :soft clips); and the SEQ field contains the sequence of bases in the read.

Read 1 shows the simplest case. The read is a forward-strand alignment, so the FLAG is 0. It aligns starting at position 5 (POS=5), with an 8-base insertion after position 7, and a 2-base deletion at positions 10-11. So, the CIGAR string is "3M8I2M2D4M", representing 3 matches, 8 insertions, 2 matches, 2 deletions, and 4 matches. We extract the sequence from the SEQ field and encode it according to the CIGAR string, as shown in Figure 3. By comparing this encoded read matrix with the reference matrix, we identify all inconsistencies, that is, potential variant locations (positions 7 and 10-11, highlighted in yellow in Figure 3).

For read 2, we observe that the reverse complement of the read matches the reference (FLAG=16), and the SEQ field of the SAM file already converts the sequence to its reverse complement. In this case, we directly encode the SEQ

Reference: AAGCTTCTCACCCTG

read1: TTCATCGATCGTCCCTG

FLAG:0 POS:5 CIGAR: 3M8I2M2D4M SEQ: TTCATCGATCGTCCCTG

| POS | 1 | 2 | 3 | 4 | 5 | 6 | 7 | 8 | 9 | 10 | 11 | 12 | 13 | 14 | 15 |
|---|---|---|---|---|---|---|---|---|---|---|---|---|---|---|---|
| REF | A | A | G | C | T | T | C | T | C | A | C | C | C | T | G |
| A | 1 | 1 | 0 | 0 | 0 | 0 | 0 | 0 | 0 | 1 | 0 | 0 | 0 | 0 | 0 |
| T | 0 | 0 | 0 | 0 | 1 | 1 | 0 | 1 | 0 | 0 | 0 | 0 | 0 | 1 | 0 |
| C | 0 | 0 | 0 | 1 | 0 | 0 | 1 | 0 | 1 | 0 | 1 | 1 | 1 | 0 | 0 |
| G | 0 | 0 | 1 | 0 | 0 | 0 | 0 | 0 | 0 | 0 | 0 | 0 | 0 | 0 | 1 |
| READ | | | | | T | T | C* | T | C | | | C | C | T | G |
| A | | | | | 0 | 0 | 2 | 0 | 0 | 0 | 0 | 0 | 0 | 0 | 0 |
| T | | | | | 1 | 1 | 2 | 1 | 0 | 0 | 0 | 0 | 0 | 1 | 0 |
| C | | | | | 0 | 0 | 3 | 0 | 1 | 0 | 0 | 1 | 1 | 0 | 0 |
| G | | | | | 0 | 0 | 2 | 0 | 0 | 0 | 0 | 0 | 0 | 0 | 1 |
| Diff | | | | | T | T | C | T | C | A | C | C | C | T | G |
| A | | | | | 0 | 0 | 2 | 0 | 0 | 1 | 0 | 0 | 0 | 0 | 0 |
| T | | | | | 0 | 0 | 2 | 0 | 0 | 0 | 0 | 0 | 0 | 0 | 0 |
| C | | | | | 0 | 0 | 2 | 0 | 0 | 0 | 1 | 0 | 0 | 0 | 0 |
| G | | | | | 0 | 0 | 2 | 0 | 0 | 0 | 0 | 0 | 0 | 0 | 0 |

Figure 3: Diagram of encoding method for read with FLAG=0. The red box shows the encoding process of CIGAR "I", the inserted bases are compressed into one column and added to the vector at the insertion position. In this example, the 8-base insertion ("8I") at a $C$ nucleotide position would generate a vector $[2\ 2\ 3\ 2]^T$, calculated as $[2\ 2\ 2\ 2]^T$ (insertion) $+[0\ 0\ 1\ 0]^T$ (insertion position base $C$). The blue box shows "D", the corresponding column is encoded as a zero vector. "Diff" represents the absolute difference between the REF matrix and the READ matrix. After subtraction, all the differences (highlighted in yellow) will be displayed.

string into the matrix, as shown in Figure 4. The encoding method for "I" and "D" in the CIGAR string is the same as when FLAG=0 (in this example, the CIGAR string only contains "D").

For reads 3 and 4, the reads are split into segments recorded as separate SAM lines. One of them is marked as primary alignment, and the additional alignments of the read are marked as supplementary alignments. To preserve the chimeric relationship inherent in split-read alignments, we encode both the primary alignment and its supplementary counterparts into a single matrix.

For read 3, the supplementary and primary alignments share the same strand orientation. Thus, the FLAG for the primary alignment is 0 (forward), and the FLAG for the supplementary alignment is 2048 (forward), as shown in Figure 2, read 3. Alternatively, the primary alignment is 16 (reverse complementary), and the supplementary alignment is 2064 (reverse complementary). Then, we position them according to relative genomic coordinates, representing the intervening regions with zeros. The encoding results of read 3 is shown in Figure 5.

For read 4, the supplementary alignments are on the opposite strand relative to the primary alignment (the FLAG for the primary alignment is 0 and for the supplementary alignment is 2064, or the FLAG for the primary alignment is 16 and for the supplementary alignment is 2048), indicating an inversion. From Figure 2, read 4, we can see that the SEQ records for these two segments are different and complement each other. If we encode them directly, the matrix will be the same as the reference matrix, as shown at the bottom of Figure 6, yielding a null difference matrix and losing the inversion information. Therefore, we use the primary direction as the benchmark and convert the SEQ string of supplementary alignments into their reverse complements. In other words, we use the original sequence of read 4 to encode the supplementary part instead of using the string recorded in the SEQ field. The middle part of Figure 6 shows our method. This preserved the inversions (highlighted in yellow) after matrix subtraction.

Reference: AAGCTTCTCACCCTG

read2: CAGGGGAATT

FLAG:16 POS:1 CIGAR: 2M2D3M3D5M SEQ: AATTCCCCTG

| POS | 1 | 2 | 3 | 4 | 5 | 6 | 7 | 8 | 9 | 10 | 11 | 12 | 13 | 14 | 15 |
|---|---|---|---|---|---|---|---|---|---|---|---|---|---|---|---|
| REF | A | A | G | C | T | T | C | T | C | A | C | C | C | T | G |
| A | 1 | 1 | 0 | 0 | 0 | 0 | 0 | 0 | 0 | 1 | 0 | 0 | 0 | 0 | 0 |
| T | 0 | 0 | 0 | 0 | 1 | 1 | 0 | 1 | 0 | 0 | 0 | 0 | 0 | 1 | 0 |
| C | 0 | 0 | 0 | 1 | 0 | 0 | 1 | 0 | 1 | 0 | 1 | 1 | 1 | 0 | 0 |
| G | 0 | 0 | 1 | 0 | 0 | 0 | 0 | 0 | 0 | 0 | 0 | 0 | 0 | 0 | 1 |
| READ | A | A | | | T | T | C | | | | C | C | C | T | G |
| A | 1 | 1 | 0 | 0 | 0 | 0 | 0 | 0 | 0 | 0 | 0 | 0 | 0 | 0 | 0 |
| T | 0 | 0 | 0 | 0 | 1 | 1 | 0 | 0 | 0 | 0 | 0 | 0 | 0 | 1 | 0 |
| C | 0 | 0 | 0 | 0 | 0 | 0 | 1 | 0 | 0 | 0 | 1 | 1 | 1 | 0 | 0 |
| G | 0 | 0 | 0 | 0 | 0 | 0 | 0 | 0 | 0 | 0 | 0 | 0 | 0 | 0 | 1 |
| Diff | A | A | G | C | T | T | C | T | C | A | C | C | C | T | G |
| A | 0 | 0 | 0 | 0 | 0 | 0 | 0 | 0 | 0 | 1 | 0 | 0 | 0 | 0 | 0 |
| T | 0 | 0 | 0 | 0 | 0 | 0 | 0 | 1 | 0 | 0 | 0 | 0 | 0 | 0 | 0 |
| C | 0 | 0 | 0 | 1 | 0 | 0 | 0 | 0 | 1 | 0 | 1 | 0 | 0 | 0 | 0 |
| G | 0 | 0 | 1 | 0 | 0 | 0 | 0 | 0 | 0 | 0 | 0 | 0 | 0 | 0 | 0 |

Figure 4: Diagram of encoding method for read with FLAG=16. The blue box shows "D", the corresponding column is encoded as a zero vector $[0\ 0\ 0\ 0]^T$. "Diff" represents the absolute difference between the REF matrix and the READ matrix. After subtraction, all the differences (highlighted in yellow) will be displayed.

Reference: AAGCTTCTCACCCTG

read3: AAGCCCCTG

Primary:

FLAG:0 POS:1 CIGAR: 4M5S SEQ: AAGCCCCTG

Supplementary:

FLAG:2048 POS:11 CIGAR: 4S5M SEQ: AAGCCCCTG

| POS | 1 | 2 | 3 | 4 | 5 | 6 | 7 | 8 | 9 | 10 | 11 | 12 | 13 | 14 | 15 |
|---|---|---|---|---|---|---|---|---|---|---|---|---|---|---|---|
| REF | A | A | G | C | T | T | C | T | C | A | C | C | C | T | G |
| A | 1 | 1 | 0 | 0 | 0 | 0 | 0 | 0 | 0 | 1 | 0 | 0 | 0 | 0 | 0 |
| T | 0 | 0 | 0 | 0 | 1 | 1 | 0 | 1 | 0 | 0 | 0 | 0 | 0 | 1 | 0 |
| C | 0 | 0 | 0 | 1 | 0 | 0 | 1 | 0 | 1 | 0 | 1 | 1 | 1 | 0 | 0 |
| G | 0 | 0 | 1 | 0 | 0 | 0 | 0 | 0 | 0 | 0 | 0 | 0 | 0 | 0 | 1 |
| READ | A | A | G | C | | | | | | | C | C | C | T | G |
| A | 1 | 1 | 0 | 0 | 0 | 0 | 0 | 0 | 0 | 0 | 0 | 0 | 0 | 0 | 0 |
| T | 0 | 0 | 0 | 0 | 0 | 0 | 0 | 0 | 0 | 0 | 0 | 0 | 0 | 1 | 0 |
| C | 0 | 0 | 0 | 1 | 0 | 0 | 0 | 0 | 0 | 0 | 1 | 1 | 1 | 0 | 0 |
| G | 0 | 0 | 1 | 0 | 0 | 0 | 0 | 0 | 0 | 0 | 0 | 0 | 0 | 0 | 1 |
| Diff | A | A | G | C | T | T | C | T | C | A | C | C | C | T | G |
| A | 0 | 0 | 0 | 0 | 0 | 0 | 0 | 0 | 0 | 1 | 0 | 0 | 0 | 0 | 0 |
| T | 0 | 0 | 0 | 0 | 1 | 1 | 0 | 1 | 0 | 0 | 0 | 0 | 0 | 0 | 0 |
| C | 0 | 0 | 0 | 0 | 0 | 0 | 1 | 0 | 1 | 0 | 0 | 0 | 0 | 0 | 0 |
| G | 0 | 0 | 0 | 0 | 0 | 0 | 0 | 0 | 0 | 0 | 0 | 0 | 0 | 0 | 0 |

Figure 5: Diagram of encoding method for reads with supplementary alignment on the same strand. The green box highlights the encoding results of primary alignment, and the purple box highlights the results of supplementary alignment. The intervening regions are encoding with zeros. After subtraction, all the differences (highlighted in yellow) will be displayed.

Reference: AAGCTTCTCACCCTG

read4: AAGCTTCTCCAGGGT

Primary:

FLAG:0    POS:1    CIGAR: 9M6S    SEQ: AAGCTTCTCCAGGGT

Supplementary:

FLAG:2064    POS:10    CIGAR: 6M9S    SEQ: ACCCTGGAGAAGCTT

| POS | 1 | 2 | 3 | 4 | 5 | 6 | 7 | 8 | 9 | 10 | 11 | 12 | 13 | 14 | 15 |
|---|---|---|---|---|---|---|---|---|---|---|---|---|---|---|---|
| REF | A | A | G | C | T | T | C | T | C | A | C | C | C | T | G |
| A | 1 | 1 | 0 | 0 | 0 | 0 | 0 | 0 | 0 | 1 | 0 | 0 | 0 | 0 | 0 |
| T | 0 | 0 | 0 | 0 | 1 | 1 | 0 | 1 | 0 | 0 | 0 | 0 | 0 | 1 | 0 |
| C | 0 | 0 | 0 | 1 | 0 | 0 | 1 | 0 | 1 | 0 | 1 | 1 | 1 | 0 | 0 |
| G | 0 | 0 | 1 | 0 | 0 | 0 | 0 | 0 | 0 | 0 | 0 | 0 | 0 | 0 | 1 |

Converting the supplements to their reverse complementary

| READ | A | A | G | C | T | T | C | T | C | C | A | G | G | G | T |
|---|---|---|---|---|---|---|---|---|---|---|---|---|---|---|---|
| A | 1 | 1 | 0 | 0 | 0 | 0 | 0 | 0 | 0 | 0 | 1 | 0 | 0 | 0 | 0 |
| T | 0 | 0 | 0 | 0 | 1 | 1 | 0 | 1 | 0 | 0 | 0 | 0 | 0 | 0 | 1 |
| C | 0 | 0 | 0 | 1 | 0 | 0 | 1 | 0 | 1 | 1 | 0 | 0 | 0 | 0 | 0 |
| G | 0 | 0 | 1 | 0 | 0 | 0 | 0 | 0 | 0 | 0 | 0 | 1 | 1 | 1 | 0 |
| Diff | A | A | G | C | T | T | C | T | C | A | C | C | C | T | G |
| A | 0 | 0 | 0 | 0 | 0 | 0 | 0 | 0 | 0 | 1 | 1 | 0 | 0 | 0 | 0 |
| T | 0 | 0 | 0 | 0 | 0 | 0 | 0 | 0 | 0 | 0 | 0 | 0 | 0 | 1 | 1 |
| C | 0 | 0 | 0 | 0 | 0 | 0 | 0 | 0 | 0 | 1 | 1 | 1 | 1 | 0 | 0 |
| G | 0 | 0 | 0 | 0 | 0 | 0 | 0 | 0 | 0 | 0 | 0 | 1 | 1 | 1 | 1 |

**Without** converting the supplements to their reverse complementary

Null difference matrix → Losing variants information

| READ | A | A | G | C | T | T | C | T | C | A | C | C | C | T | G |
|---|---|---|---|---|---|---|---|---|---|---|---|---|---|---|---|
| A | 1 | 1 | 0 | 0 | 0 | 0 | 0 | 0 | 0 | 1 | 0 | 0 | 0 | 0 | 0 |
| T | 0 | 0 | 0 | 0 | 1 | 1 | 0 | 1 | 0 | 0 | 0 | 0 | 0 | 1 | 0 |
| C | 0 | 0 | 0 | 1 | 0 | 0 | 1 | 0 | 1 | 0 | 1 | 1 | 1 | 0 | 0 |
| G | 0 | 0 | 1 | 0 | 0 | 0 | 0 | 0 | 0 | 0 | 0 | 0 | 0 | 0 | 1 |
| Diff | A | A | G | C | T | T | C | T | C | A | C | C | C | T | G |
| A | 0 | 0 | 0 | 0 | 0 | 0 | 0 | 0 | 0 | 0 | 0 | 0 | 0 | 0 | 0 |
| T | 0 | 0 | 0 | 0 | 0 | 0 | 0 | 0 | 0 | 0 | 0 | 0 | 0 | 0 | 0 |
| C | 0 | 0 | 0 | 0 | 0 | 0 | 0 | 0 | 0 | 0 | 0 | 0 | 0 | 0 | 0 |
| G | 0 | 0 | 0 | 0 | 0 | 0 | 0 | 0 | 0 | 0 | 0 | 0 | 0 | 0 | 0 |

Figure 6: Diagram of encoding method for reads with supplementary alignment on the different strand. The green box shows the encoding results of primary alignment, and the purple box shows the results of supplementary alignment. Here, we compare the encoding results of employing reverse-complement sequence vs. native-strand sequence for supplementary SEQ when the supplementary alignment resides on the complementary strand relative to the primary alignment. Without converting the supplements to their reverse complement, we can see that the Diff matrix is null and the variant information is missing.

#### 2. Supplementary Table

The 28 genes in the Breast cancer dataset are listed in Supplementary Table 1.

**Supplementary Table 1**
The 28 genes of focus related to breast cancer.

| Chrom | Start | End | Gene | Length |
|---|---|---|---|---|
| chr2 | 211,375,717 | 212,538,802 | ERBB4 | 1,163,085 |
| chr2 | 214,725,645 | 214,809,711 | BARD1 | 84,066 |
| chr3 | 179,148,114 | 179,240,093 | PIK3CA | 91,979 |
| chr5 | 132,556,924 | 132,644,621 | RAD50 | 87,697 |
| chr6 | 151,654,148 | 152,129,604 | ESR1 | 475,456 |
| chr7 | 55,019,017 | 55,211,628 | EGFR | 192,611 |
| chr8 | 31,033,262 | 31,175,871 | WRN | 142,609 |
| chr8 | 89,933,336 | 89,984,724 | NBS1 | 51,388 |
| chr8 | 144,511,284 | 144,517,828 | RECQL4 | 6,544 |
| chr9 | 95,099,054 | 95,317,730 | FANCC | 218,676 |
| chr11 | 94,415,570 | 94,512,701 | MRE11 | 97,131 |
| chr11 | 101,029,624 | 101,130,681 | PGR | 101,057 |
| chr11 | 108,222,484 | 108,369,099 | ATM | 146,615 |
| chr12 | 56,080,025 | 56,103,507 | ERBB3 | 23,482 |
| chr13 | 32,315,480 | 32,399,672 | BRCA2 | 84,192 |
| chr14 | 67,819,779 | 68,683,118 | RAD51B | 863,339 |
| chr14 | 64,226,712 | 64,338,631 | ESR2 | 111,919 |
| chr15 | 90,717,327 | 90,816,166 | BLM | 98,839 |
| chr16 | 23,603,162 | 23,641,357 | PALB2 | 38,195 |
| chr16 | 68,737,290 | 68,835,542 | CDH1 | 98,252 |
| chr17 | 31,094,927 | 31,377,677 | NF1 | 282,750 |
| chr17 | 35,099,792 | 35,119,869 | RAD51D | 20,077 |
| chr17 | 39,688,084 | 39,728,662 | ERBB2 | 40,578 |
| chr17 | 43,044,295 | 43,125,483 | BRCA1 | 81,188 |
| chr17 | 58,692,140 | 58,735,611 | RAD51C | 43,471 |
| chr17 | 61,679,186 | 61,864,120 | BRIP1 | 184,934 |
| chr17 | 75,626,845 | 75,667,202 | RECQL5 | 40,357 |
| chr22 | 28,687,743 | 28,741,866 | CHEK2 | 54,123 |

**Supplementary Table 2**
SVs detected in in-house breast cancer data only by *g*SV.

| Chrom | Start | Gene | Length | Type | Other Information |
|---|---|---|---|---|---|
| chr2 | 211689991 | ERBB4 | 170 | DUP | |
| chr2 | 211944066 | ERBB4 | 148 | DUP | |
| chr2 | 212191391 | ERBB4 | 214 | DUP | |
| chr7 | 55167022 | EGFR | 754 | DUP | Other tools detect this DUP as an INS. Panel (c) of Figure 5 in main text shows the detail. |
| chr7 | 55176724 | EGFR | 246 | DUP | |
| chr14 | 68037106 | RAD51B | 84 | DEL | |
| chr15 | 90804992 | BLM | 935 | DEL | |
| chr17 | 61787317 | BRIP1 | 57 | DUP | Other tools detect this DUP as an INS. |

#### 3. Supplementary Figures

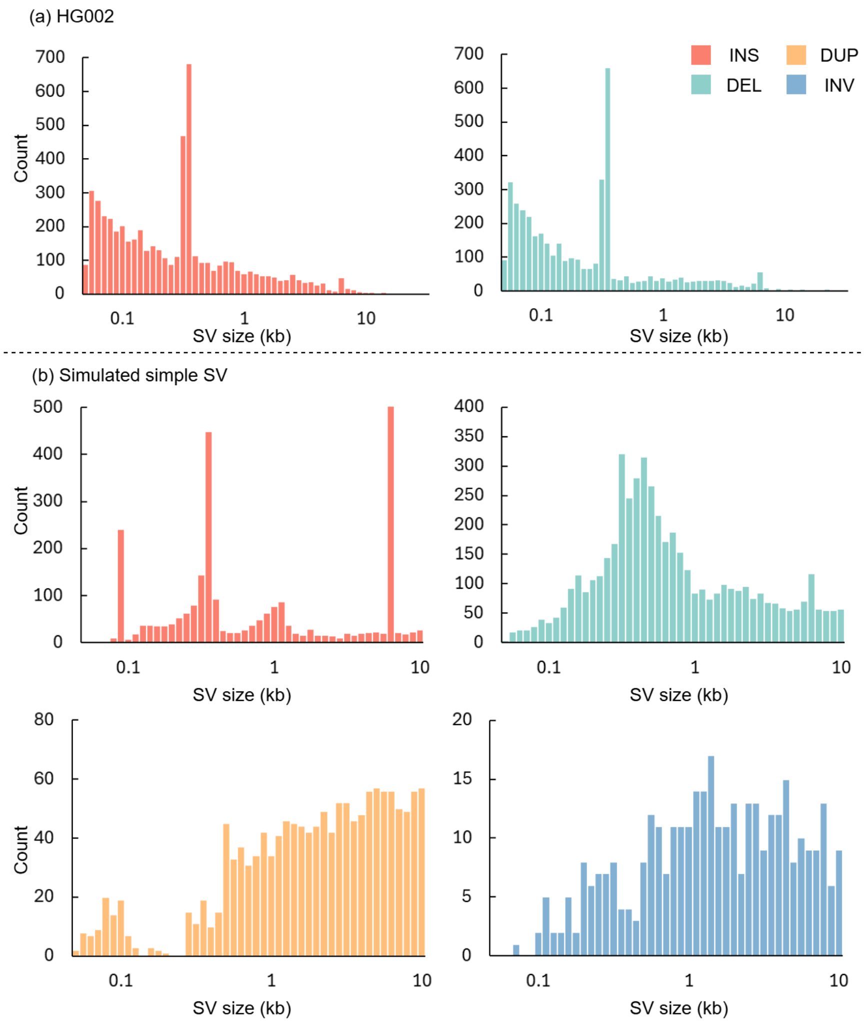


Supplementary Figure 1: The size distributions of simple SVs. The horizontal axis represents the SV length in kilobases, while the vertical axis shows the number of SVs at each length.

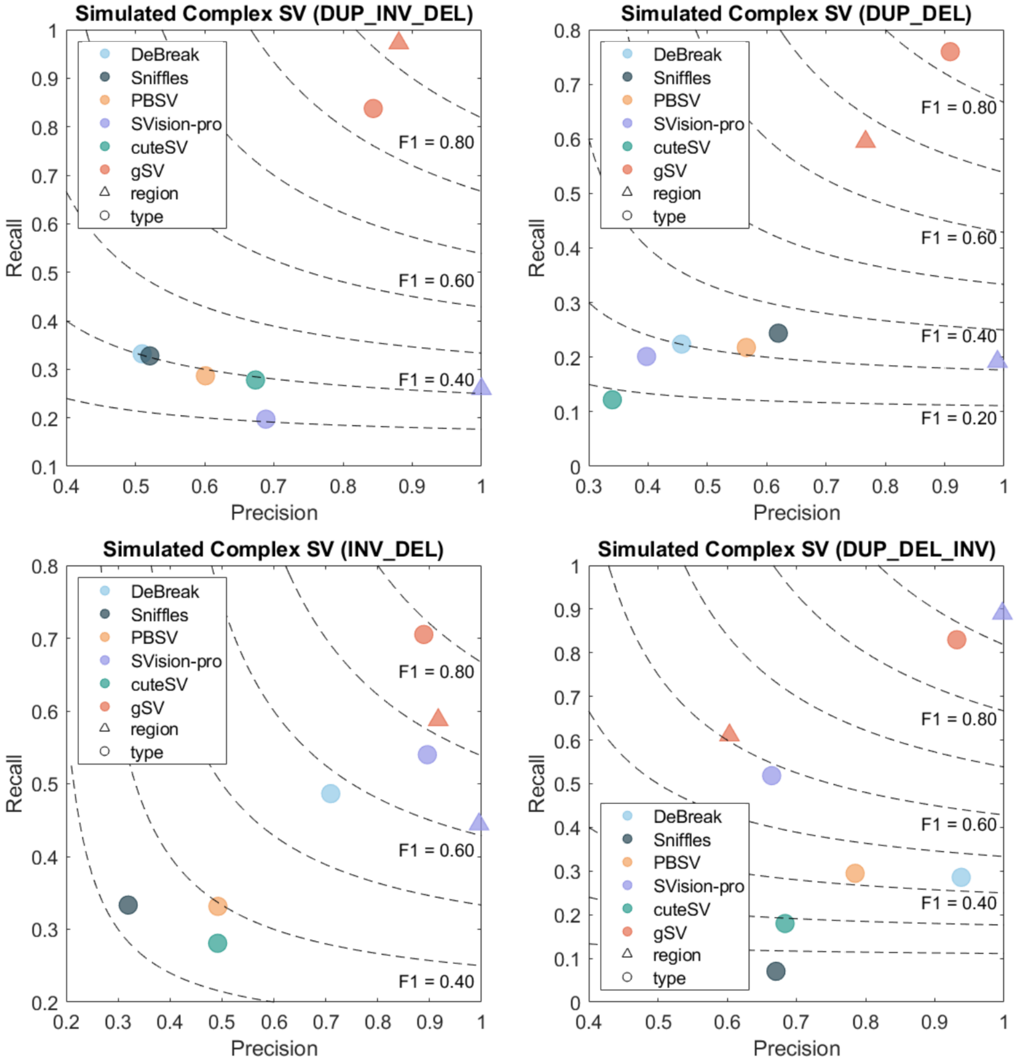


Supplementary Figure 2: The performance of different methods in detecting different types of complex SVs by evaluating the region and type, respectively. Four out of five existing tools (DeBreak, Sniffles, PBSV, and cuteSV) without complex SV detection capability perform worse than *g*SV in identifying such composite events involving multiple breakpoints. The fifth existing tool SVision-pro performs better than *g*SV in identifying region loations for DUP_DEL_INV type of SVs. But *g*SV achieves more accurate type detection than SVision-pro.

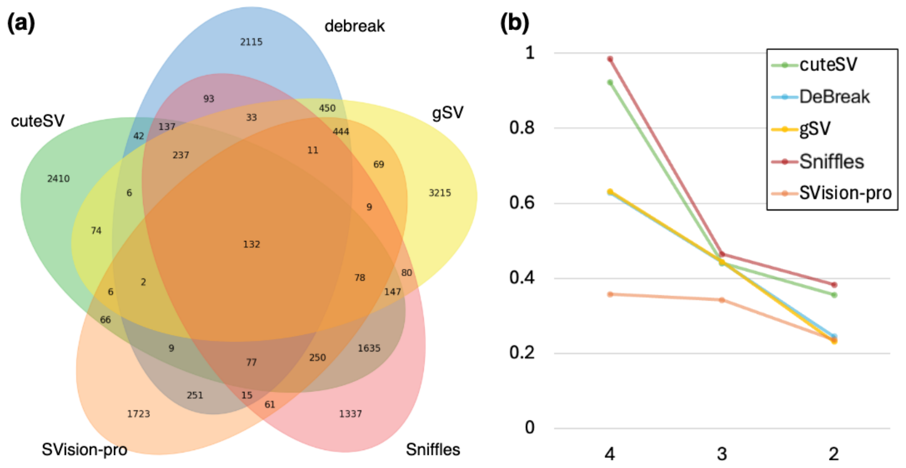


Supplementary Figure 3: Panel (a) shows a Venn diagram of the numbers of somatic SVs detected by different tools on HCC1395. Panel (b) reports the ratios of five tools in SV call sets of different confidence levels. The horizontal axis denotes the number of other tools that have co-detected each SV set. SVs co-detected by four other tools have a higher confidence level than those detected by only three other tools. It can be seen that the curves for all five tools exhibited descending trends as confidence levels decreased, indicating that all tools report more SVs for input sets with higher signal-to-noise ratio (i.e., higher percent of true SVs). Note that the relative height comparison between different lines is meaningless because we do not know the correct ratio of signals within each set, but steeper deceasing means the corresponding tool is more hesitating to report SVs for input set containing less percent of true SVs. Thus gSV, indicated by the constantly sharp decreasing from 4 to 3 and from 3 to 2, may be the most immune tool against noise (i.e., true negatives). From this logic, we argue that the 3215 SVs only reported by gSV may be largely true SVs.

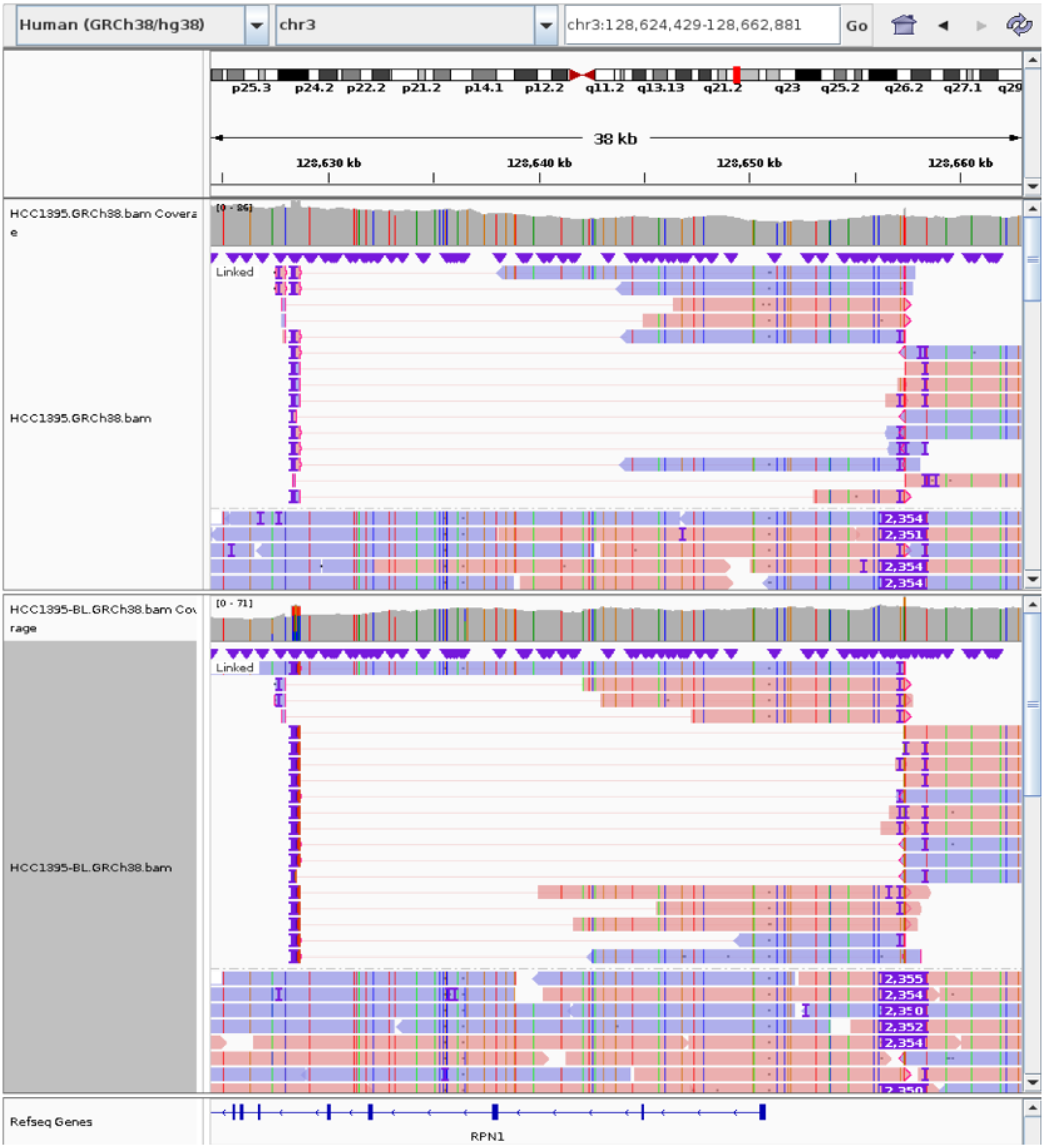


Supplementary Figure 4: A germline INV uniquely detected by *g*SV in HCC1395 and HCC1954. The INV is located within the RPN1 exonic region. Previous research [11] has demonstrated that RPN1 promotes the proliferation, migration, and invasion of breast cancer cells through the PI3K/AKT/mTOR signaling pathway.

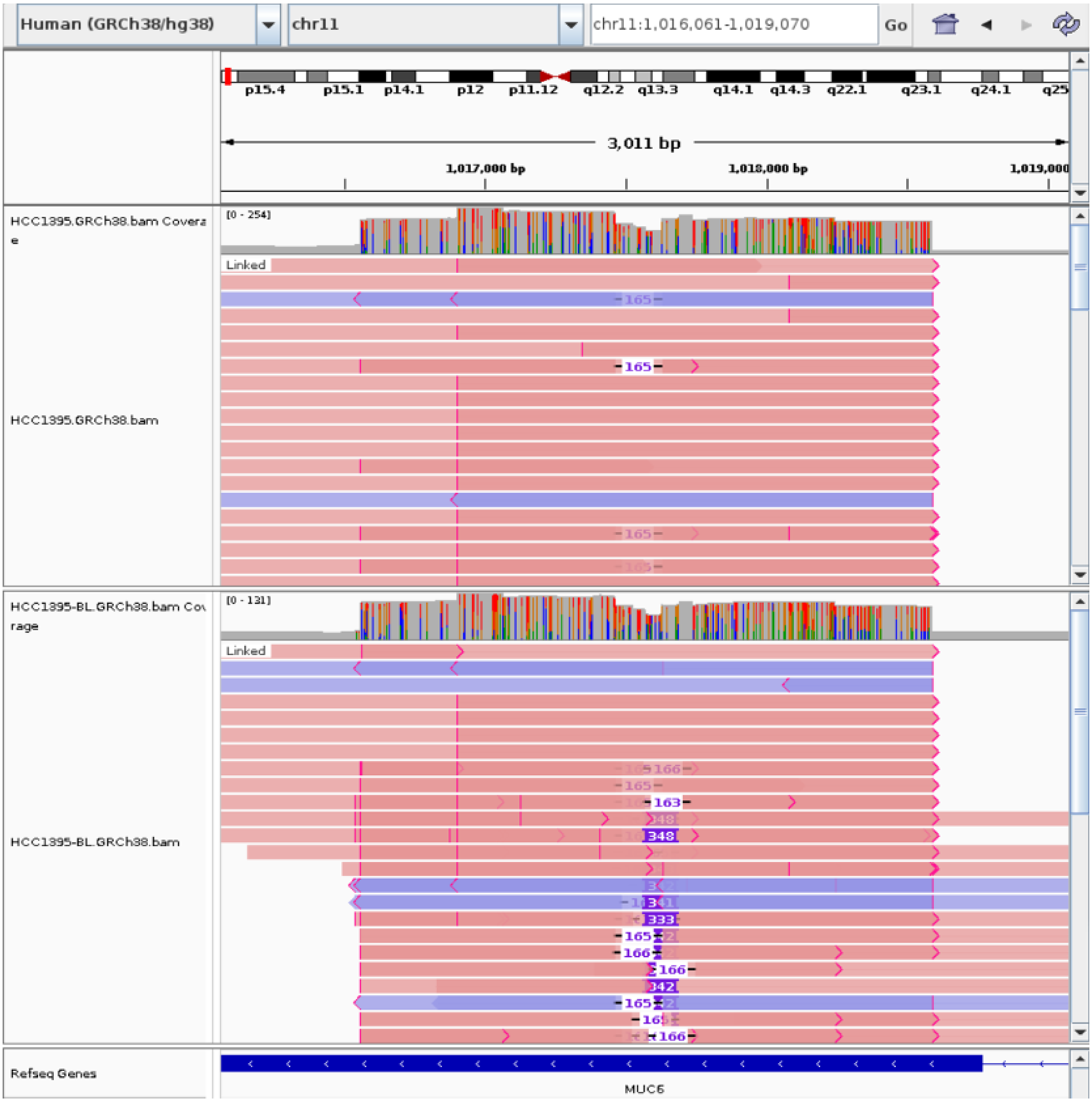


Supplementary Figure 5: Two overlapping DUPs in the exonic region of MUC6 uniquely detected by *g*SV in HCC1395. Two-step signals in the IGV visualization confirm the findings, whereas competing tools (cuteSV, DeBreak, Sniffles) detected only the second DUP, and SVision-pro failed to identify either. MUC6 is one of five secreted gel-forming mucins (MUC2, MUC5AC, MUC5B, MUC6, and MUC19) expressed from a gene cluster at chromosome 11p15. Literature review reveals that MUC6 alterations have been linked to tumor aggressiveness in cancers [12], and it is overexpressed in breast cancer [13].

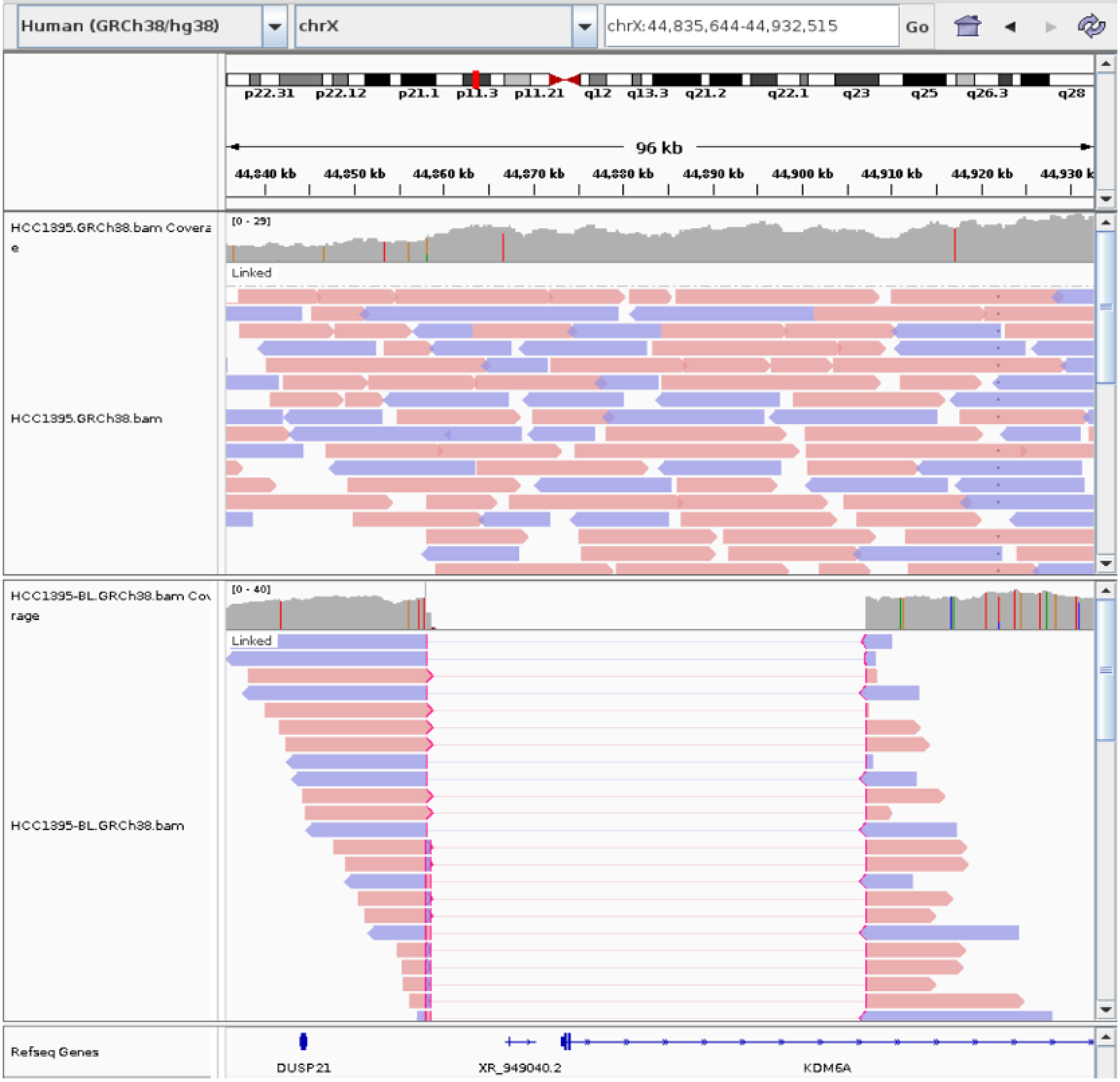


Supplementary Figure 6: A DEL uniquely detected by *g*SV in HCC1395. The DEL is located within the KDM6A exonic region. KDM6A is commonly mutated in multiple cancer types, including gastric, urothelial, pancreatic, and breast cancer. Previous research [14] has shown that silencing KDM6A promoted cell migration and transformation, demonstrated by the formation of tumor-like acini in three-dimensional culture.

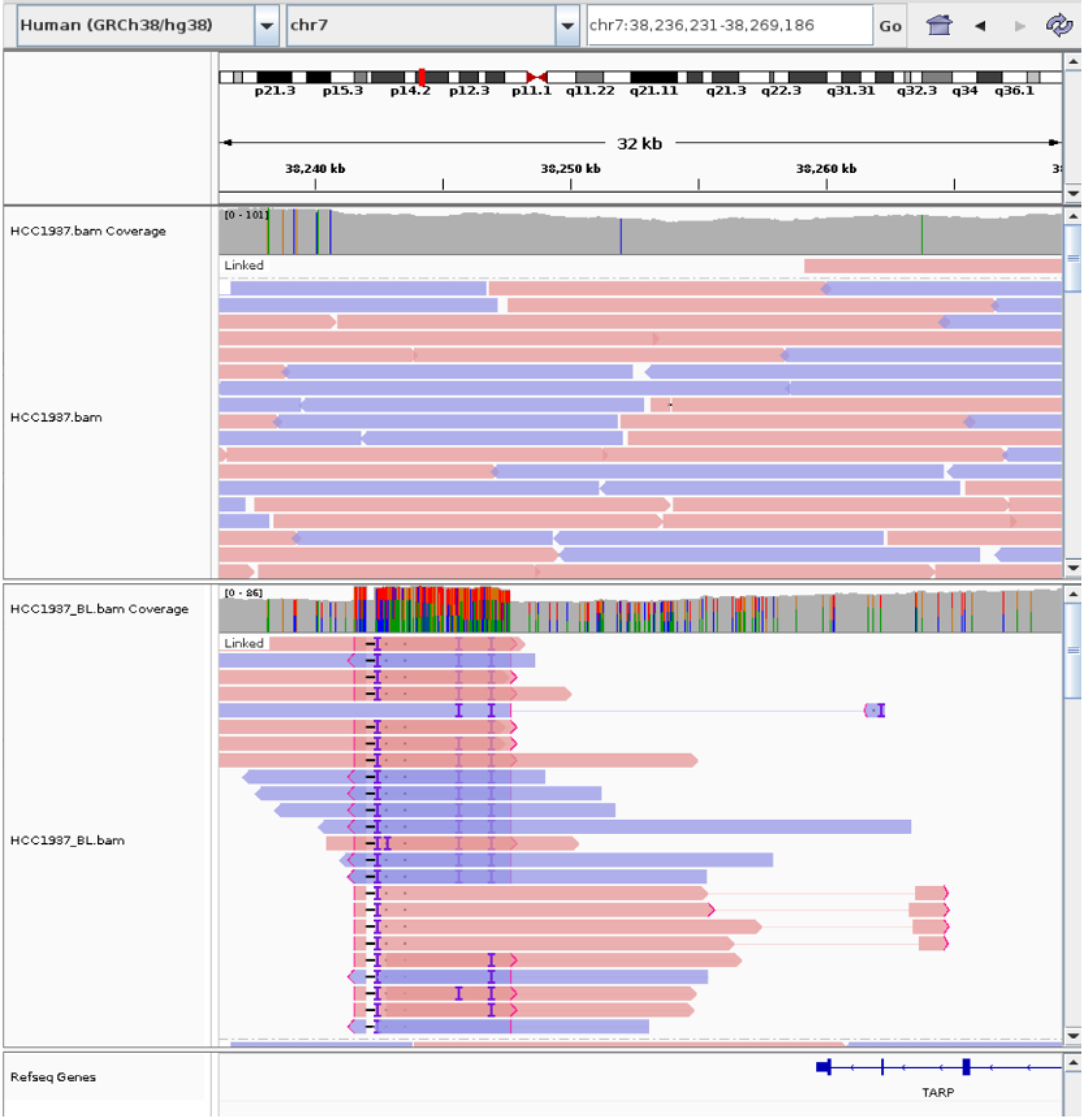


Supplementary Figure 7: A DUP uniquely detected by *g*SV in HCC1937. The DUP is located within the exonic region of the TCR gamma alternate reading frame protein (TARP, a nuclear protein expressed in prostate and breast cancer cells [15]).

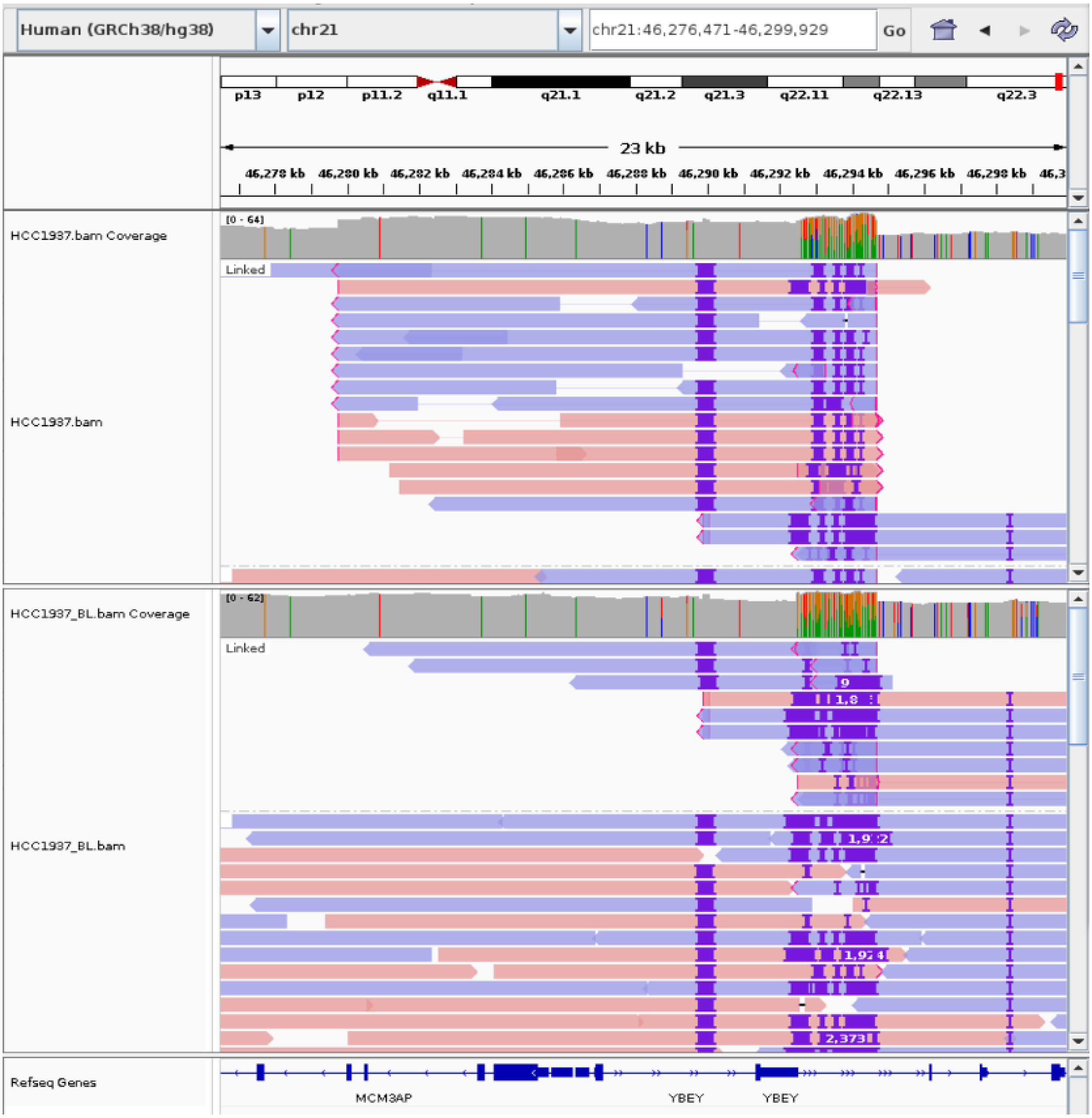


Supplementary Figure 8: A DUP uniquely detected by *g*SV in HCC1937. The DUP is located within the MCM3AP exonic region. Previous studies have shown that the lncRNA MCM3AP-AS1 may be a novel breast cancer lncRNA with high expression levels in breast cancer patients' tissue [16]. It can promote breast cancer progression via modulating the miR-28-5p/CENPF axis [17].

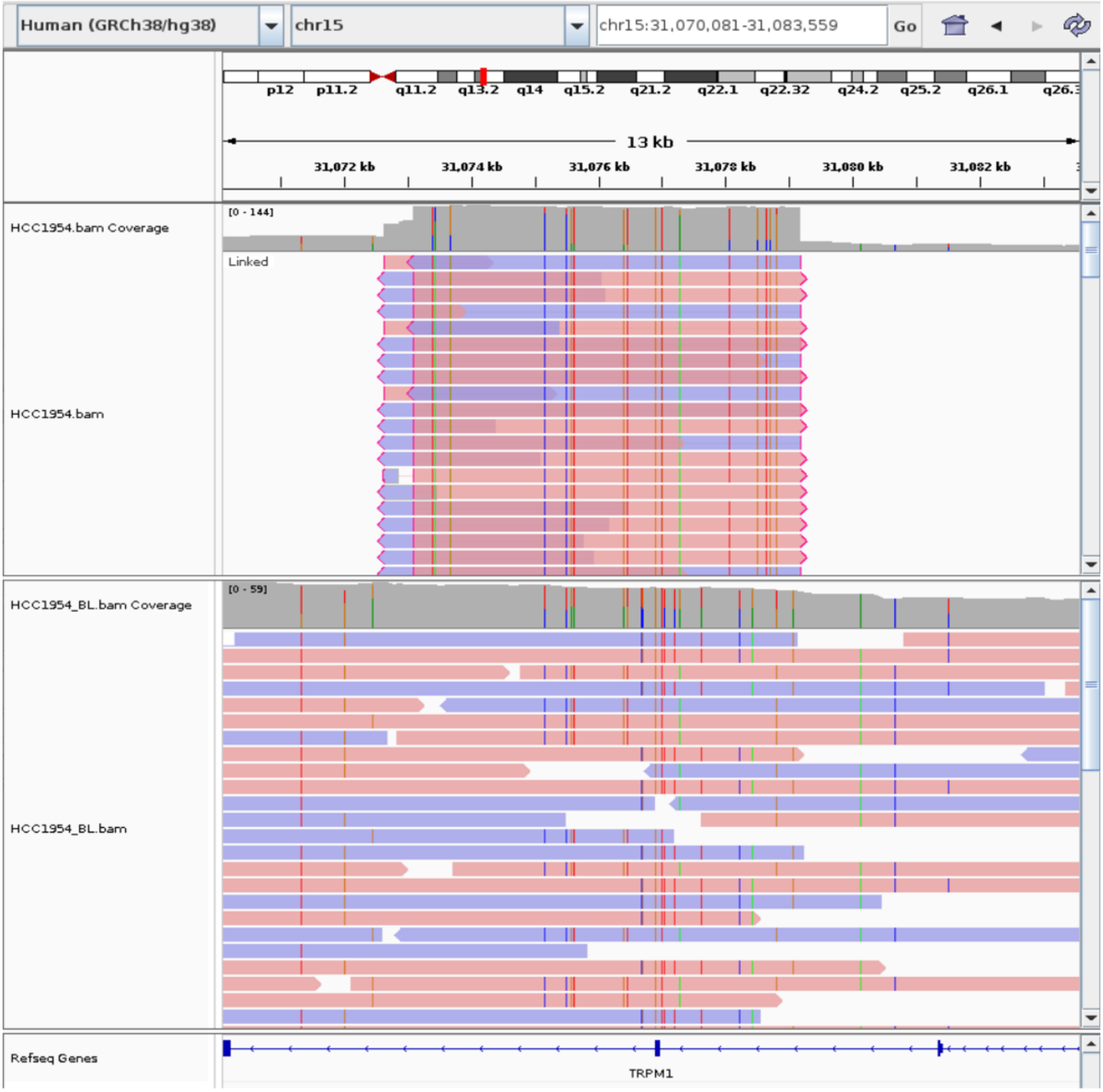


Supplementary Figure 9: A somatic complex SV (DUP+invDUP) uniquely detected by *g*SV in HCC1954. The complex SV is located within the exonic region of TRPM1. TRPM1 is a specific gene for breast carcinoma [18] and can be used as a great diagnostic tool, especially for Triple-negative Breast Cancer (TNBC).

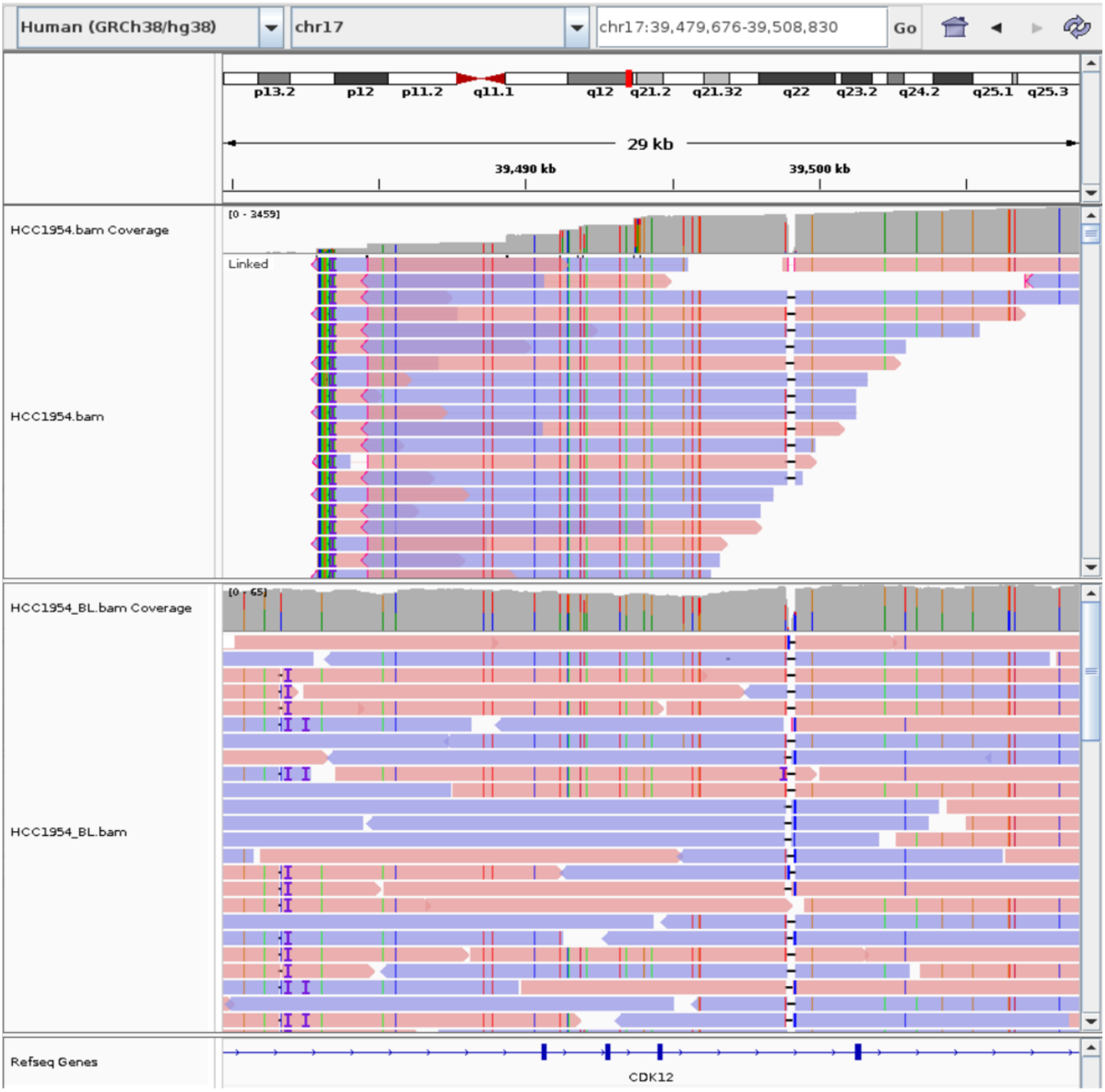


Supplementary Figure 10: A somatic complex SV (DUP+invDUP) uniquely detected by *g*SV in HCC1954. The complex SV is located within the exonic region of CDK12, which is a primary oncogene. When CDK12 is overexpressed in the normal mammary epithelium, it leads to the formation of spontaneous breast tumors, in addition to actively cooperating in chemical carcinogen- or oncogene-induced breast tumorigenesis [19].

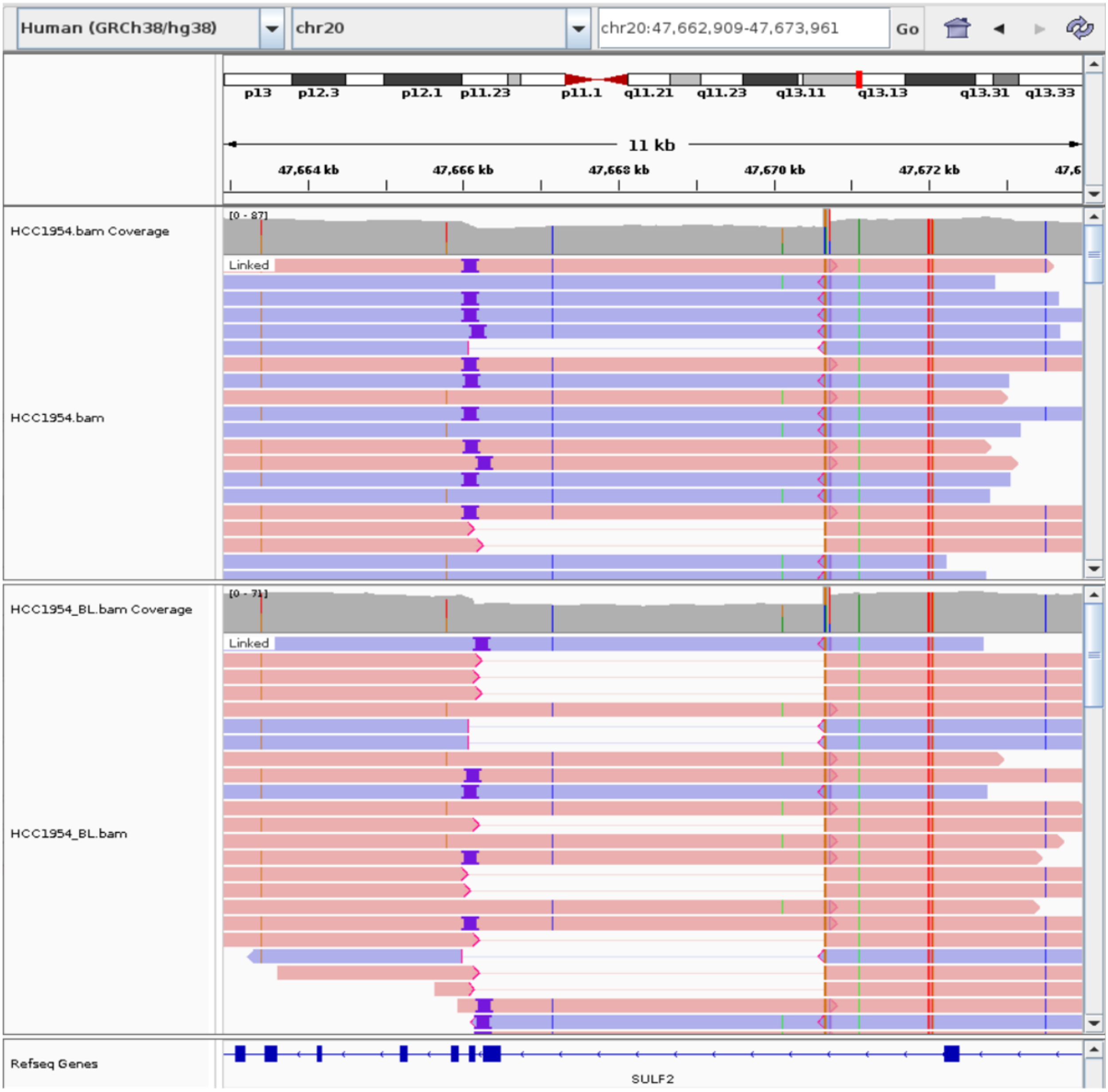


Supplementary Figure 11: A DEL uniquely detected by *g*SV in HCC1954. The DEL is located within the SULF2 exonic region. Previous research [20] has shown that SULF2 increased breast cancer proliferation, invasion, mobility, and adhesion both in vitro and in vivo.

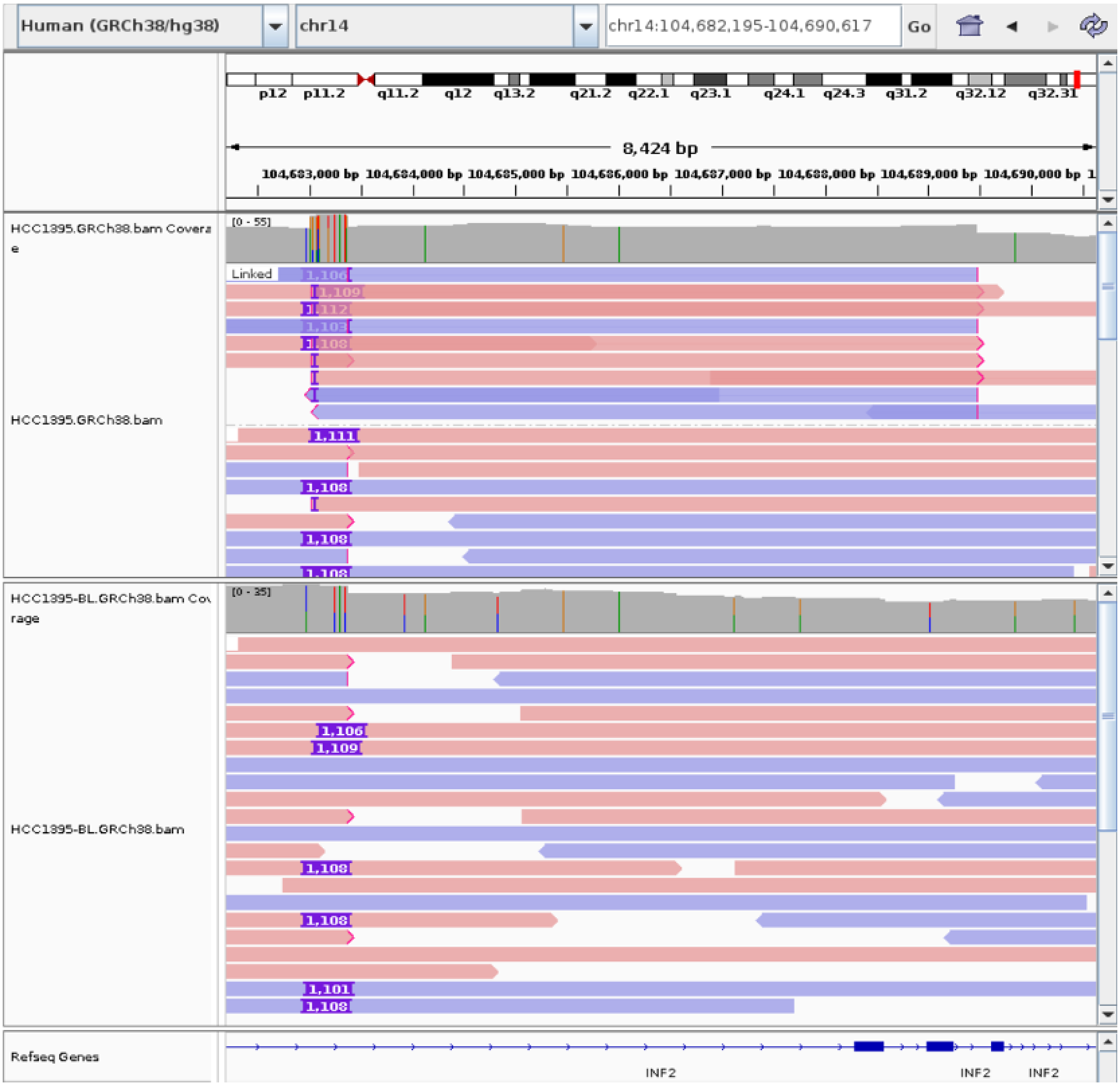


Supplementary Figure 12: A DUP uniquely detected by *g*SV in HCC1395. The DUP is located within the upstream regulatory region of INF2. Previous research [21] has shown that basal-like breast cancers frequently overexpress formin proteins FHOD1 and INF2.

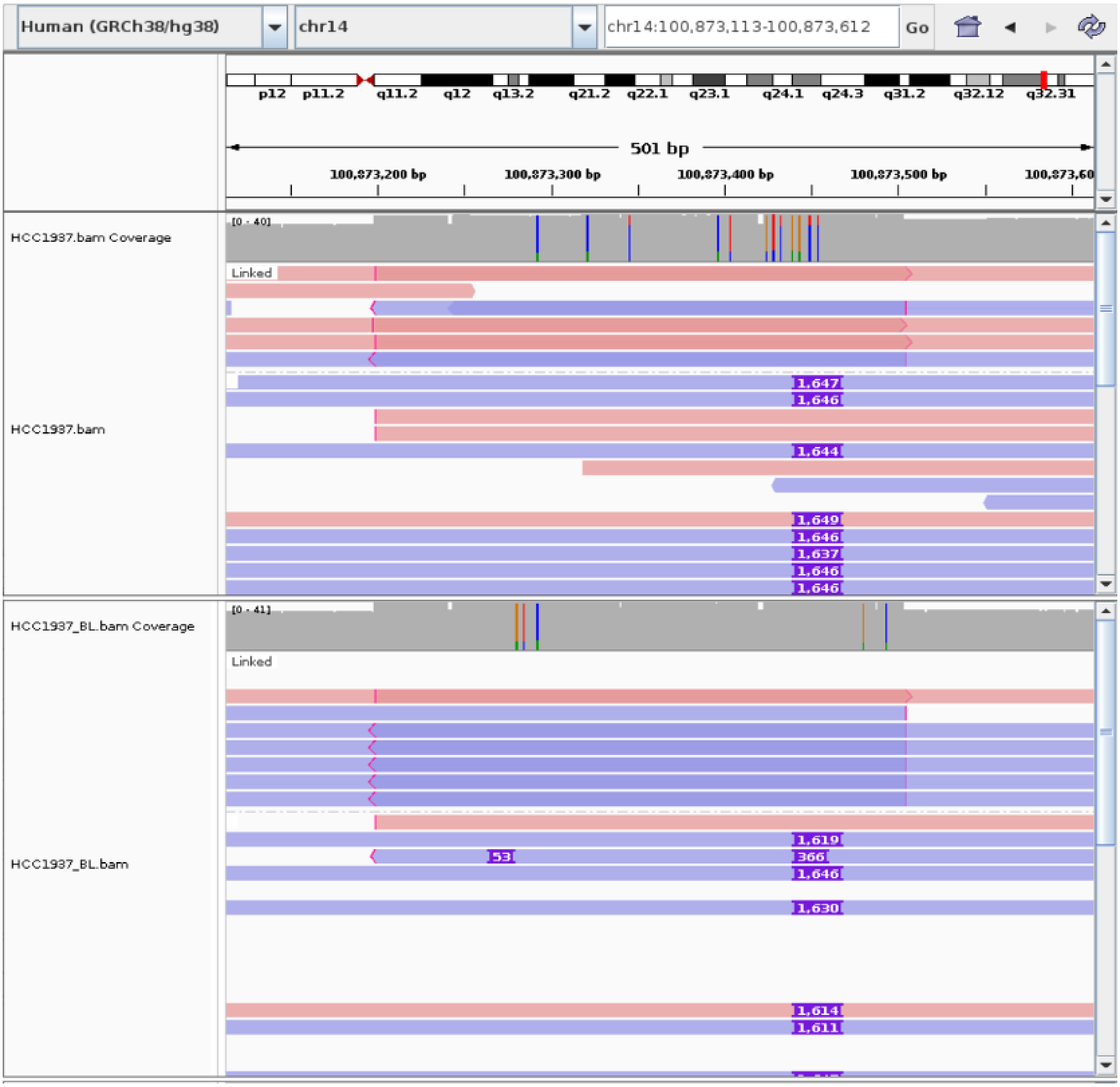


Supplementary Figure 13: A DUP uniquely detected by *g*SV in HCC1937. The DUP is located within the upstream regulatory region of MIR337. Previous research [22] has shown that this microRNA can suppress breast cancer cell migration and invasion via downregulation of ESRP1.

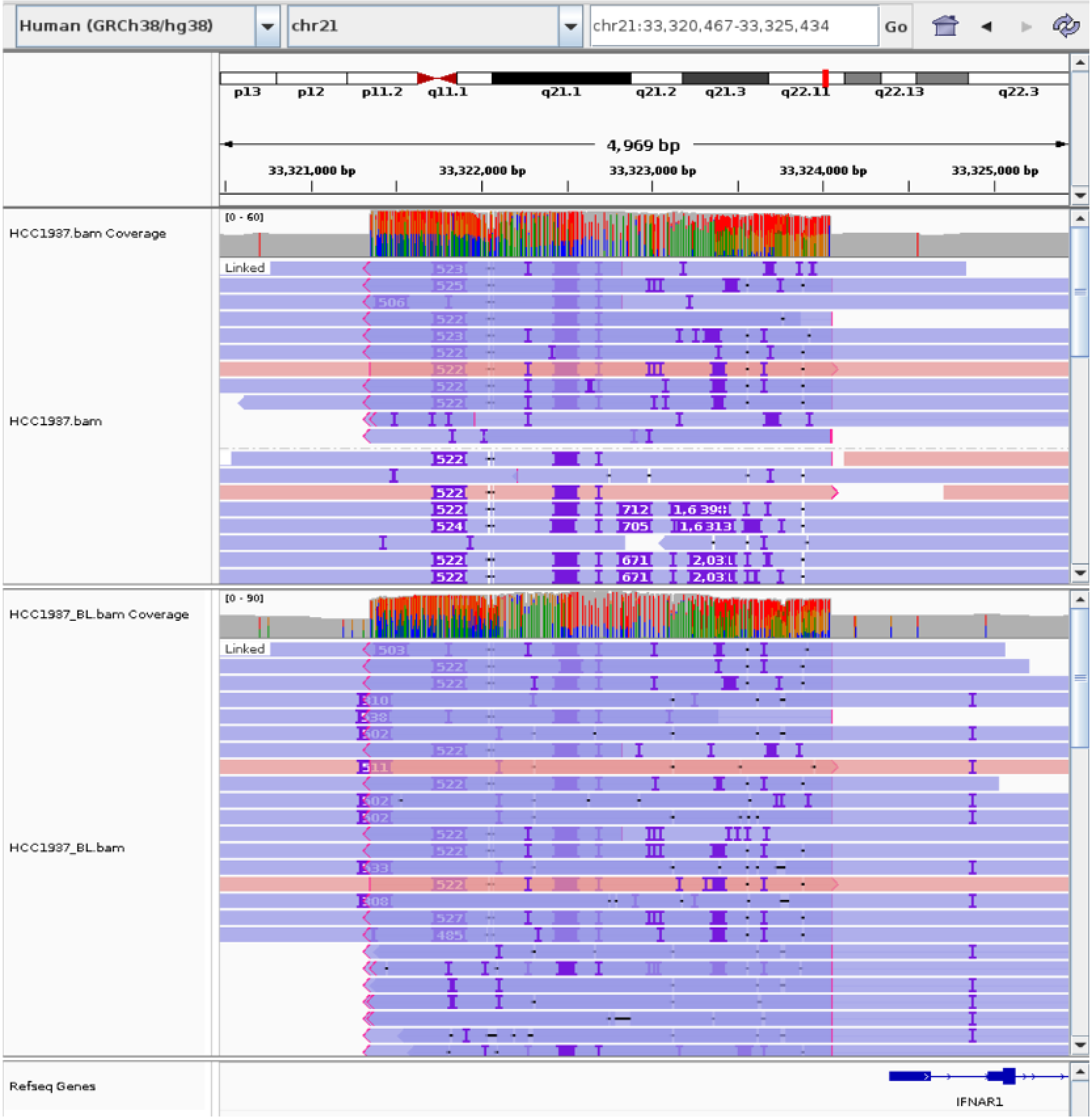


Supplementary Figure 14: A DUP uniquely detected by *g*SV in HCC1937. The DUP is located within the upstream regulatory region of IFNAR1. Previous research [23] has shown that high IFNAR1 levels in breast cancer cells are associated with poor prognosis.

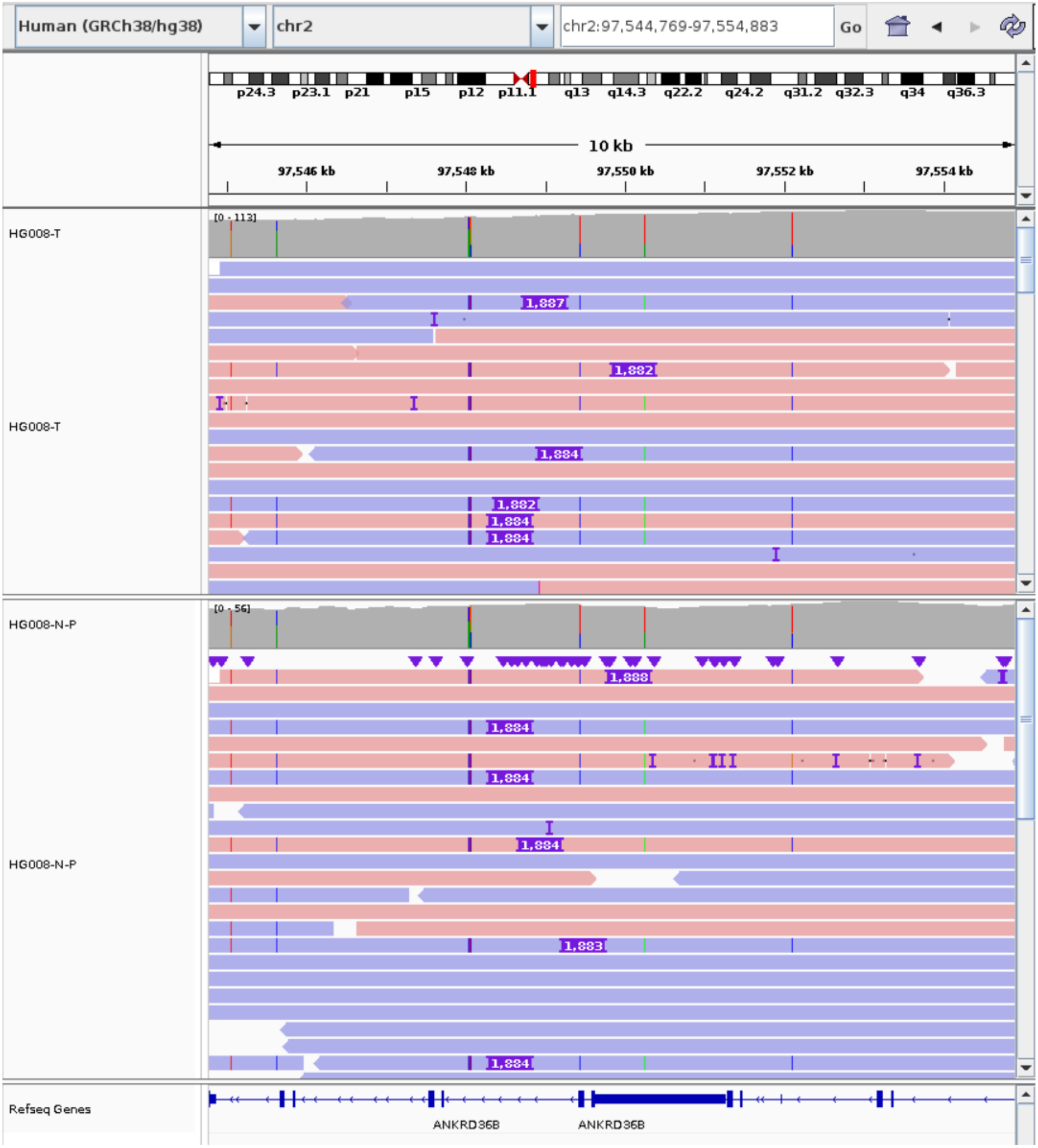


Supplementary Figure 15: A DUP uniquely detected by *g*SV in HG008. The DUP is located within the exonic region of ANKRD36B. The role of ANKRD36B in cancer is less well-established, but the Human Protein Atlas suggests it's expressed in various cancer tissues.

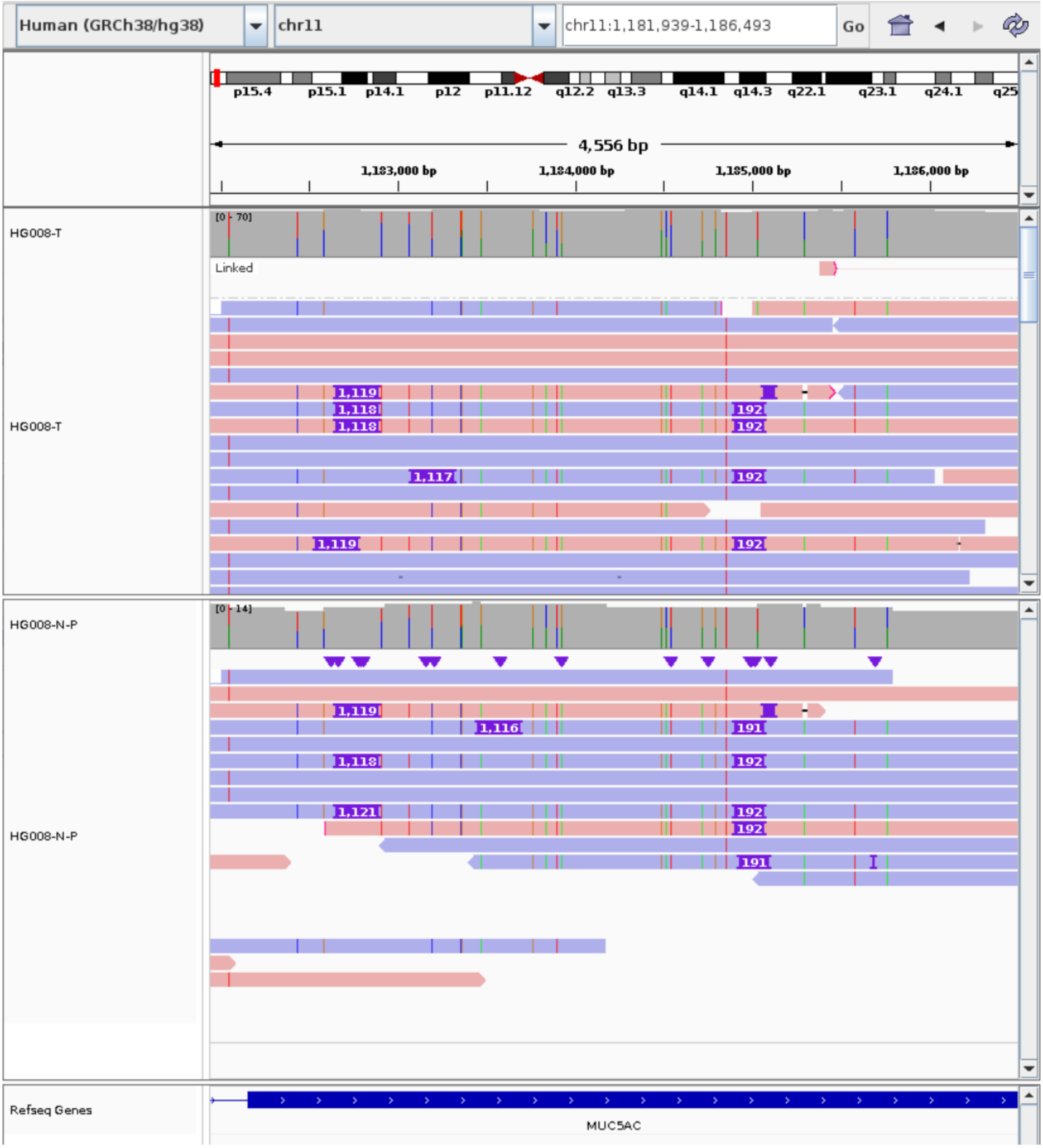


Supplementary Figure 16: A DUP uniquely detected by *g*SV in HG008. The DUP is located within the exonic region of MUC5AC. Previous research [24] has shown that MUC5AC is a secretory mucin aberrantly expressed in various cancers, e.g. lung cancer, but the relationship with breast cancer is unclear.

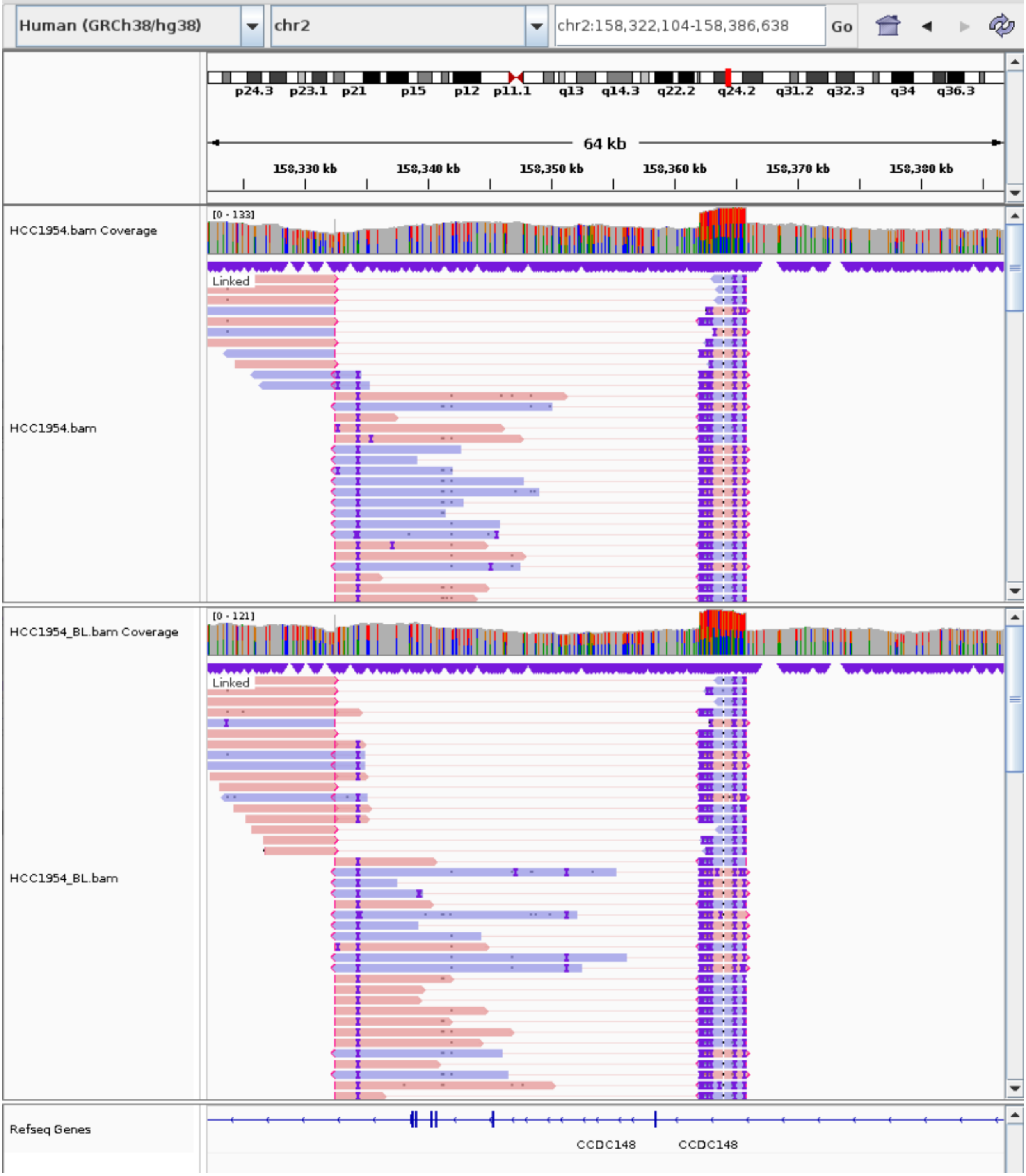


Supplementary Figure 17: An INV uniquely detected by *g*SV in HCC1954. The INV is located within the exonic region of CCDC148. Previous research [25] has found that CCDC148 gene variants are associated with the risk of aromatase inhibitors-induced musculoskeletal syndrome (AIMSS) in HR+ breast cancer patients treated with aromatase inhibitors. Further research is needed to discover and validate genetic predictors of AIMSS.
